# The transcription factor OsbZIP48 governs rice responses to zinc deficiency

**DOI:** 10.1101/2023.05.18.541251

**Authors:** Shubao Hu, Binbin Du, Guangmao Mu, Zichen Jiang, Hui Li, Yuxinrui Song, Baolei Zhang, Jixing Xia, Hatem Rouached, Luqing Zheng

**Affiliations:** College of Life Sciences, Nanjing Agricultural University, Nanjing 210095, China; State Key Laboratory for Conservation and Utilization of Subtropical Agro-bioresources, College of Life Science and Technology, Guangxi University, Nanning 530005, China; Department of Plant, Soil, and Microbial Sciences, Plant Resilience Institute, Michigan State University, East Lansing, MI 48824

**Keywords:** rice, Zn deficiency, transcription factors, OsbZIP48, zinc deficiency response element (ZDRE), zinc transporter genes

## Abstract

Zinc deficiency is the most prevalent micronutrient disorder in rice and leads to delayed development and decreased yield. Nevertheless, despite its primary importance, how rice responds to zinc deficiency remains poorly understood. Herein, we present genetic evidence that OsbZIP48 is essential for regulating rice responses to zinc deficiency. Using the reverse genetics approach, genetic inactivation of *OsbZIP48* in rice seedlings caused a hyper sensitivity to zinc deficiency, associated with a significant decrease in the root-to-shoot translocation of zinc. Consistently, *OsbZIP48* was constitutively expressed in roots, slightly induced by zinc deficiency in shoots, and localized into nuclei induced by Zn deficiency. Comparative transcriptome analysis of the wild-type plants and *osbzip48* mutant grown under zinc deficiency enabled the identification of OsbZIP48 target genes, including key zinc transporter genes (*OsZIP4* and *OsZIP8*). We demonstrated that OsbZIP48 controlled the expressions of these genes by directly binding to their promoters, specifically to the zinc deficiency response element (ZDRE) motif. Collectively, we showed that the *OsbZIP48* gene encodes for a transcription factor in rice, and demonstrates its critical role in the response to zinc deficiency in this crop. This knowledge is crucial for the design of rice plants that are resilient to the globally prevalent zinc limitation through zinc bio-fortification programs.

## Introduction

Zinc (Zn) is an essential micronutrient that plays key roles in all organisms. It is a non-redox active element that acts as a cofactor for a wide range of proteins that are involved in many central biochemical pathways, including carbohydrate, protein, nucleic-acid, and lipid metabolisms (Broadley et al., 2012). Zn-containing proteins also play vital roles in DNA replication and in the regulation of gene expression. Zn deficiency can severely affect both the growth and development of plants. Typical symptoms of Zn deficiency in plants are stunted growth and the “little leaf” phenotype, often combined with chlorosis. In contrast, excessive Zn uptake can be toxic to plants. Zn toxicity can be observed in mining areas and with low pH soil conditions, which inhibits root elongation and causes chlorosis of young leaves (Broadley et al., 2012). Therefore, Zn homeostasis in plants is tightly regulated (Sinclair & Krämer, 2012).

The regulation of Zn homeostasis relies on a suite of genes that encode proteins involved in Zn uptake, translocation, storage, and distribution (Palmer & Guerinot, 2009; Sinclair & Krämer, 2012; Huang et al., 2022). Zn deficiency causes an up-regulation of the expression of many Zn-homeostasis related genes, such as ZRT/IRT-like protein (ZIP) family transporters encoding genes. In the model plant *Arabidopsis*, the two transcription factors (TFs) bZIP19 and bZIP23 (both members of F-bZIP family), were identified as important in the mediation of the plant response to Zn deficiency. These TFs were identified via yeast one hybrid (Y1H) screening with the promoter of *ZIP4*, a gene whose expression was strongly induced by Zn deficiency in *Arabidopsis* (Assunção et al., 2010). The expression of *bZIP19* and *bZIP23* are up-regulated by Zn deficiency at the transcriptional level and they mediate the up-regulation of *ZIP* genes containing a binding site for Zn Deficiency Response Elements (ZDREs) in their promoters (Assunção et al., 2010; Inaba et al., 2015). *In planta*, the *bzip19bzip23* double mutant is hypersensitive to Zn deficiency and accumulates less Zn in its roots and shoots (Assunção et al., 2010). Further investigation showed that bZIP19 and bZIP23 function as Zn sensor by its Zn^2+^ binding activity of the cysteine (Cys) and histidine (His)-rich motif and coordinate downstream Zn deficiency responses (Lilay et al., 2021). Through a heterologous complementation assay, the four F-bZIP TFs; TabZIPF1-7DL (TabZIP180) and TabZIPF4-7AL (TabZIP55) from wheat, and HvbZIP56 and HvbZIP62 from barley, were found to partially rescue the Zn-dependent growth phenotype of the *Arabidopsis bzip19bzip23* double mutant (Evens et al., 2017; Nazri et al., 2017). In addition, ZDREs were found in the promoters of several wheat and barley *ZIP* genes, and TabZIP180 was shown to bind to ZDRE in the *TaZIP* promoter. However, their expression patterns and subcellular localizations differ from *Arabidopsis* F-bZIP proteins in barley and wheat (Evens et al., 2017; Nazri et al., 2017). For example, the expression of *TabZIPF1* and *TabZIPF4* from wheat were up-regulated by Zn deficiency, while two barley genes *HvbZIP56* and *HvbZIP62* showed little change at early stage of Zn deficiency and the expression of *HvbZIP56* in the shoot was down-regulated after long-term Zn starvation; bZIP19 and bZIP23 are localized in the nucleus, whereas the HvbZIP56-GFP signal was observed in both nucleus and cytosol when expressed in tobacco cells (Inaba et al., 2015; Evens et al., 2017; Nazri et al., 2017). Despite the seemingly conserved function of F-bZIP TFs across plants in response to Zn deficiency, the exact physiological function of F-bZIP proteins and their downstream regulating genes awaits further examinations. Therefore, further investigation is needed as the result may hold unexpected specificity for the response to Zn limitation of each species.

Rice (*Oryza sativa* L.) is a model grass and crop system, and also serves as a staple food for more than half of the world’s population. Zn deficiency is one of the most prevalence micronutrient limitation that significantly affects rice yield and quality (Broadley et al., 2012). However, the regulatory mechanisms in response to Zn deficiency in rice has not been well understood. The transport activity of Zn was confirmed for OsZIP1, OsZIP3, OsZIP4, OsZIP5, OsZIP7, OsZIP7a, OsZIP8, and OsZIP9 either in yeast *zrt1/zrt2* mutants or via transgenic analyses of model plant systems (Ramesh, 2003; Ishimaru et al., 2005; Yang et al., 2009; Lee et al., 2010a; Lee et al., 2010b; Sasaki et al., 2015; Ricachenevsky et al., 2018). Among these, the expressions of *OsZIP4*, *OsZIP5*, and *OsZIP8* were induced in response to Zn deficiency, and their overexpressing in rice significantly affected the distribution of Zn between roots and shoots, suggesting that the three ZIP transporter are involved in root-to-shoot Zn translocation and distribution (Ishimaru et al., 2005; Lee et al., 2010a; Lee et al., 2010b). Recently, OsZIP3 and OsZIP4 were demonstrated to be involved in delivering Zn to new developing tissues in rice based on detailed expression pattern analysis and mutant analysis (Sasaki et al., 2015; Mu et al., 2021). On the other hand, studies showed that OsZIP9 mainly contribute to Zn uptake from soil in rice roots under Zn-limiting condition (Huang et al., 2020; Tan et al., 2020; Yang et al., 2020). Moreover, another study showed that the rice F-bZIP TF OsbZIP48 (LOC_Os06g50310), which has high similarity in amino acid sequences with bZIP19 and bZIP23, could bind to a ZDRE-containing *AtZIP4* promoter fragment in a Y1H assay and complement the *Arabidopsis* mutants *bzip19bzip23* as a heterologous system (Lilay et al., 2020). This raises an important question on the biological function of OsbZIP48 and its direct downstream genes in rice. The present study comprehensively characterized *OsbZIP48* in rice at the functional level. By combining a variety of approaches we revealed the essential role of OsbZIP48 in regulating rice Zn transport, rice growth and development in Zn limiting conditions, and further provide evidences for the specificity of OsbZIP48 role in rice compared to dicots (*i.e.* Arabidopsis).

## Results

### Expression analysis of *OsbZIP48*

The temporal and spatial expression pattern of *OsbZIP48* was examined using rice samples harvested from the field. *OsbZIP48* was almost constitutively expressed at all organs (Fig. 1A). At the vegetative growth stage, stronger expression was found in the leaf blade and sheath (Fig. 1A). We also investigated the effect of metal deficiencies on the expression of *OsbZIP48*. Three marker genes (*OsZIP4*, *OsIRT1* and *OsCOPT5*) for Zn, Fe, and Cu deficiency, respectively, were used to evaluate the status of the deficiency. *OsZIP4* (Ishimaru et al., 2005) was induced by more than 10 times in the Zn-deficient roots, and *OsIRT1* (Bughio et al., 2002) and *OsCOPT5* (Yuan et al., 2010) were induced by 6.9 and 6.5 times, respectively, in the Fe-and Cu-deficient roots (Supplemental Fig. S1). Under different metal deficiencies, the expression of *OsbZIP48* was slightly induced by Zn-deficiency in the shoots, but not in the roots, and its expression was not affected by Fe-, Mn-or Cu-deficiency (Fig. 1B). To further gain insight into the response of *OsbZIP48* to changes in Zn availability in rice, the expression profile of this TF was analyzed in roots and shoots of WT plants grown in either Zn deficiency or Zn sufficiency conditions for 9 d. The Zn deficiency marker gene *OsZIP4* was significantly upregulated after 3 and 5 d of Zn deficiency in the root and shoot, respectively, and further induced thereafter (Fig. 1C). The *OsbZIP48* transcript remained unchanged in roots during the 9 d of Zn deficiency application (Fig. 1D), and increased slightly in shoots after 7 d of Zn deficiency (Fig. 1D).

**Figure 1.**
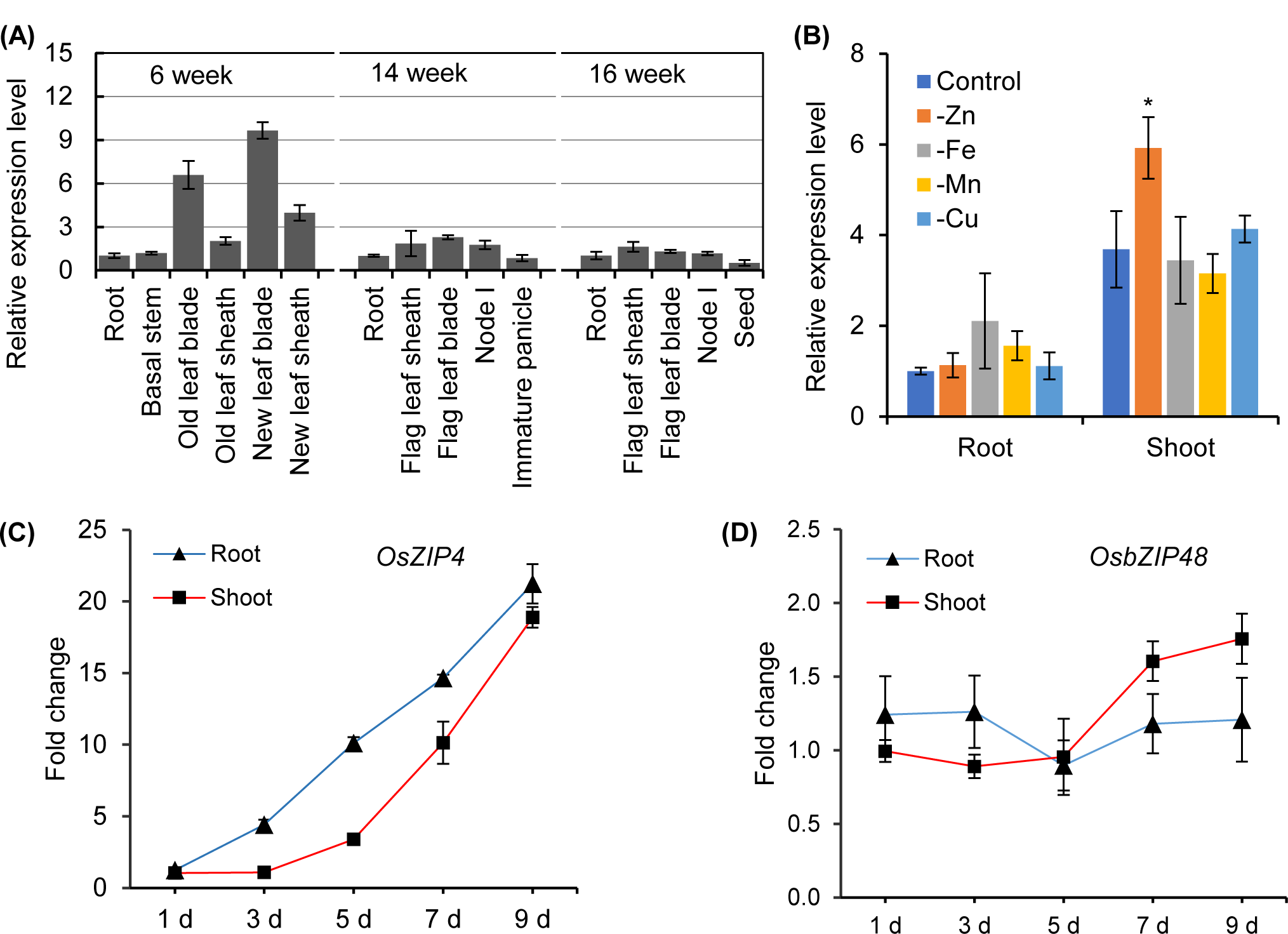
Expression patterns of *OsbZIP48*. (A) The expression level of *OsbZIP48* in various tissues across different growth stages and tissues. Different tissues were sampled from rice grown in a paddy field. Gene expression was determined by RT-qPCR. (B) The relative expression level of *OsbZIP48* in roots and shoots under deficiencies of various metal elements. Two-week-old rice seedlings were grown in nutrient solutions without Zn, Fe, Mn, and Cu, respectively, for 7 d. The relative expression levels of *OsZIP4* (C) and *OsbZIP48* (D) in roots and shoots of 2-week-old rice seedlings grown in -Zn solutions for 9 d. Expression relative to roots at 6 weeks (A), roots under control conditions (B), and the root under +Zn at 1 d (C and D) are shown. *Actin* was used as an internal standard. Data are means ± SD (n = 3). Statistical comparison between control and treatment was performed by student t-test. Asterisks indicate significant differences from the control (*P < 0.05).

### OsbZIP48 is a transcriptional activator

OsbZIP48 was predicted to encode a putative TF. To examine whether OsbZIP48 would have transcriptional activation activity, a yeast assay was employed. The result indicated that pGBKT7-bZIP48 but not the negative control vector pGBKT7 could rescue the yeast cells growth on the selective growth medium (Supplemental Fig. S2), this result supports the role of OsbZIP48 as transcriptional activator in rice, and raises the question for its physiological role *in planta* in response to -Zn.

### *OsbZIP48* genetic inactivation causes hypersensitivity to Zn limitation in rice

To investigate the physiological role of OsbZIP48 in rice, *osbzip48* knockout mutants were generated via CRISPR/Cas9. Two independent homozygous mutant lines (*bzip48-1* and *bzip48-2*; Supplemental Fig. S3) were identified by sequencing. *bzip48-1* contains a 57 bp deletion in target gRNA1, causing a loss of 19 amino acid in the conserved N-terminal of the F-bZIP transcription factor; while *bzip48-2* contains a 1 bp insertion in target gRNA2, producing a frameshift mutation and a premature stop codon (Supplemental Fig. S3). The growth capacity of WT plants and *osbzip48* knockout mutants grown under different Zn regimes, were compared. In the presence of Zn, the growth performances of the two *osbzip48* mutants were similar to that of WT (Supplemental Fig. S4A and S5). However, the *osbzip48* mutants gradually showed a severe growth inhibition under Zn deficiency compared with WT (Supplemental Fig. S4 and S5). After 15 d of Zn deficiency, a significantly less root biomass and severe necrosis in leaf blades and leaf sheaths were observed in *osbzip48* mutants compared with WT (Supplemental Fig. S4, B-D and S5). After a further 10 d of Zn deficiency stress, *osbzip48* mutants showed major necrosis and had around 44.0% and 67.5% less biomass than controls (Fig. 2, A-E), which ultimately stopped plant growth. To determine the specificity of OsbZIP48 in Zn deficiency response in rice, we treated WT plants and *osbzip48* mutants with Fe-, Cu-, or Mn-deficient solutions for 25 d, both *osbzip48* mutants and WT plants displayed similar phenotypes (Supplemental Fig. S6).

**Figure 2.**
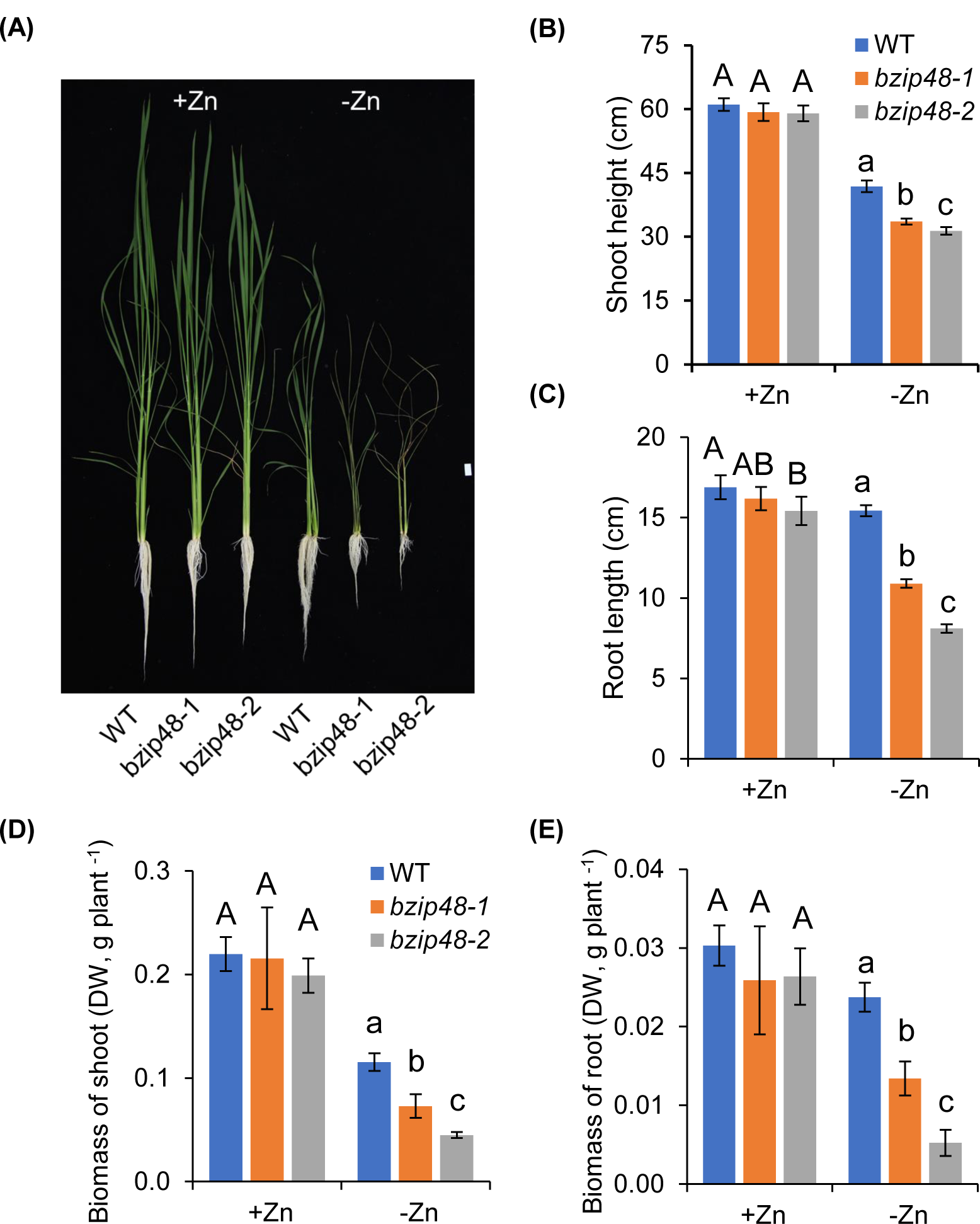
*osbzip48* mutants are hypersensitive to -Zn. **(A)** Phenotype of WT plants and two *osbzip48* mutant lines (*bzip48-1* and *bzip48-2*) under Zn-sufficient (+Zn) or Zn-deficient (-Zn) conditions for 25 d in 1/2 Kimura B solution. Shoot height (B), root length (C), and biomass of shoot (D) and root (E) were shown. Significant differences between WT plants and the two *osbzip48* mutant lines were calculated by one-way ANOVA followed by Tukey’s multiple comparison test. Data are means ± SD (n = 5). Different letters indicate significant differences (P<0.05). Scale bars = 3 cm.

To assess whether the necrosis phenotype under Zn-deficiency observed in the mutants was indeed caused by genetic inactivation of *OsbZIP48*, we performed a genetic complementation assay by introducing the genomic sequence of *OsbZIP48* (*ProbZIP48::bZIP48-GFP*) into the mutant *bzip48-1*. Analysis with two independent complementary lines showed that the hypersensitivity phenotypes to Zn-deficiency were rescued (Fig. 3, A-E). These results indicate that *osbzip48* mutants are hypersensitive to Zn deficiency, and that the assessed phenotypes are specific to Zn deficiency. OsbZIP48 appears to be required for rice survival under long-term Zn deficiency conditions.

**Figure 3.**
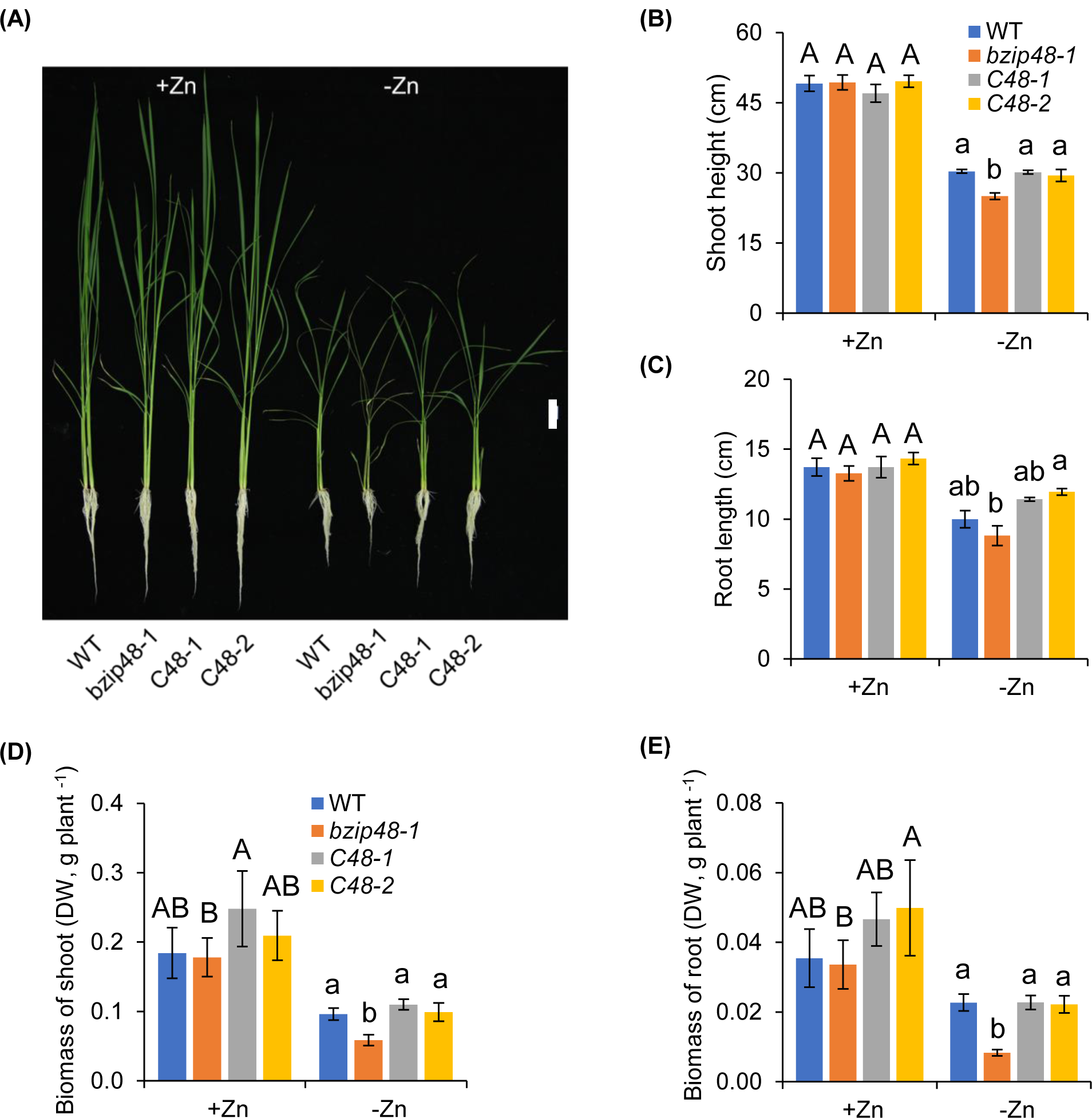
Genetic complementation of *osbzip48* mutant. (A) Phenotype of WT, *bzip48-1* mutant, and two complementary lines (*C48-1*, *C48-2*) under Zn-sufficient (+Zn) or Zn-deficient (-Zn) conditions for 28 d in 1/2 Kimura B solution. Shoot height (B), root length (C), and biomass of shoot (D) and root (E) were shown. Significant differences between WT plants and *bzip48-1* mutant, or *bzip48-1* mutant and *C48-1*, *C48-2* lines were calculated by one-way ANOVA followed by Tukey’s multiple comparison test. Data are means ± SD (n = 5). Different letters indicate significant differences (P<0.05). Scale bars = 3 cm.

### Cellular and subcellular localization of OsbZIP48

Then we used the GFP-fused OsbZIP48 complementation lines to examine the cell specificity localization of OsbZIP48 by immunostaining with a GFP antibody. In the roots, the signal was observed almost in the whole roots with a strong expression in the root tips (Fig. 4, A-C). No signal was detected in the roots of wild-type rice (Fig. 4, D-F), indicating the specificity of the antibody. In the shoots, OsbZIP48 was highly expressed in the shoot axillary meristem, shoot apical meristem (Fig. 4, G-J). We also observed that OsbZIP48 was higher expressed in the root vascular bundles based on the cross-section immunostaining (Fig. 4, K-N).

**Figure 4.**
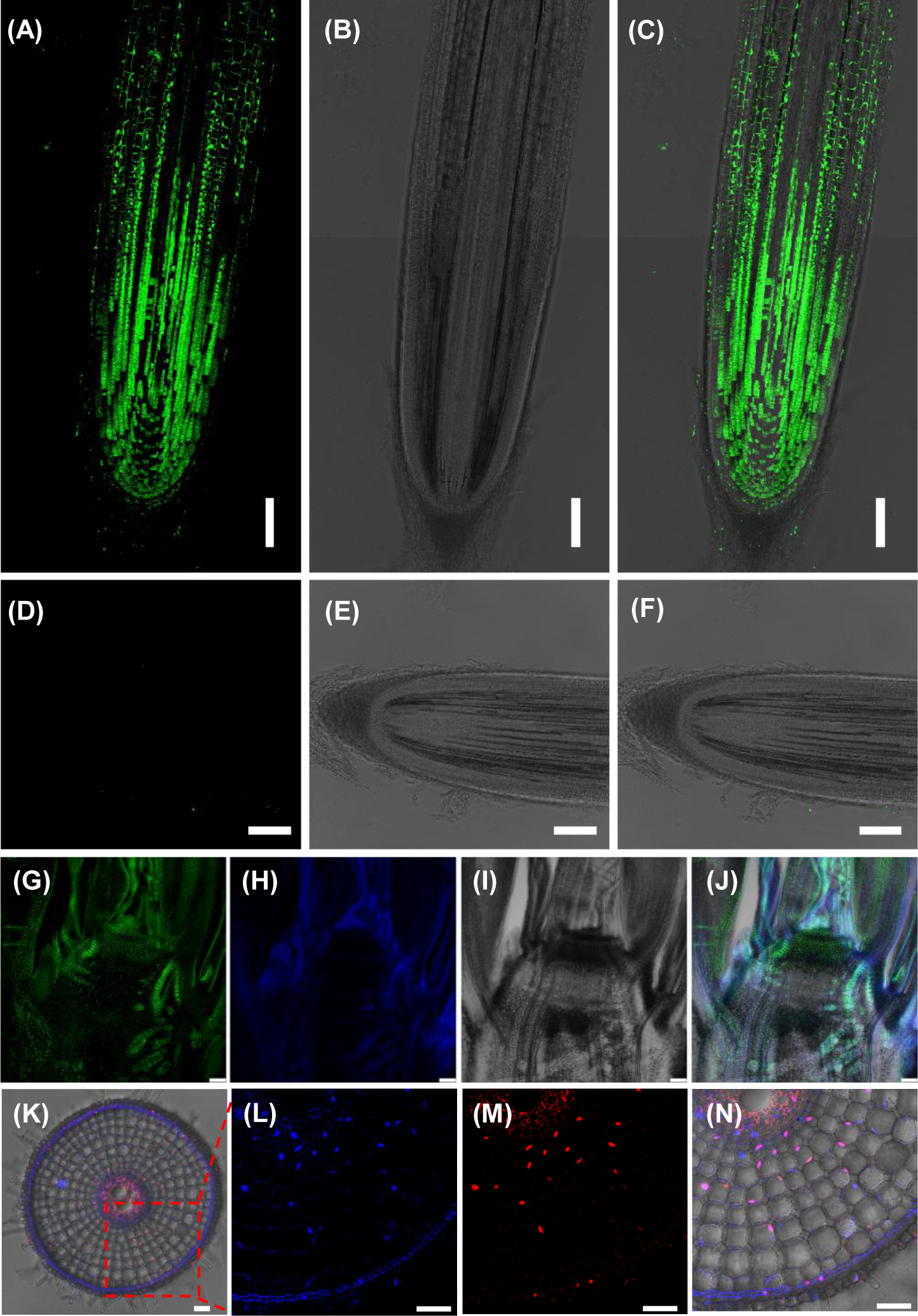
Tissue and subcellular localization of OsbZIP48. Immunohistochemical staining of OsbZIP48 with an anti-green fluorescent protein (GFP) antibody was performed in the roots (A-C, and K-N) and shoots basal node (G-J) of *ProbZIP48*::*bZIP48*-GFP transgenic rice, and roots (D-F) of wild-type rice as a negative control. Green (A-J) and red (K-N) correspond to antibody signal, cyan correspond to DAPI signal. The signal of antibody (A, G and M), the transmitted light image (B, E, I), autofluorescence of cell wall and nuclei stained by DAPI (H and L), and the merged image (C, F, J, K and N) are shown. Scale bars are 100 μm.

OsbZIP48 was reported to be localized to the nucleus and also in the cytosol when expressed in the Arabidopsis *bzip19bzip23* double mutant (Lilay et al., 2020). To examine whether OsbZIP48 protein shows similar subcellular localization in rice plants, immunostaining co-stained with 4’,6-diamidino-2-phenylindole (DAPI) for nuclei was performed in GFP-fused OsbZIP48 rice roots. Results showed that OsbZIP48 is localized in both nuclei and cytosol of the rice cells (Fig. 4, K-N). Given that OsbZIP48 is a transcription factor involved in Zn deficiency responses, we wondered whether Zn deficiency had an impact on the subcellular localization of OsbZIP48, we observed the GFP fluorescent signal in the GFP-fused OsbZIP48 rice roots under both Zn deficiency and Zn-sufficient conditions, as shown in Supplemental Fig. S7, OsbZIP48 proteins were detected both in the nuclei and cytosol, and Zn deficiency significantly promote it localized to nuclei.

### OsbZIP48 affects Zn root-to-shoot translocation and distribution in rice in -Zn conditions

The expression pattern of *OsbZIP48* and the phenotypes of its genetic inactivation under Zn deficiency led to the proposition of a general role of it in the regulation of Zn homeostasis in rice. Compared with the role of F-bZIP TFs, its plausible involvement in the regulation of Zn transport in rice was tested first, independent of Zn availability. In either presence or absence of Zn, no difference in Zn accumulation was observed in roots or shoots in *osbzip48* mutants compared with WT plants (Supplemental Fig. S8, A and B). Noteworthy, Zn concentrations were significantly lower in both young leaf blades and leaf sheaths of *osbzip48* mutants compared with WT plants under Zn deficiency (Supplemental Fig. S9, A and B). However, no difference was found in Zn concentrations of old tissues between *osbzip48* mutants and WT plants under the same conditions (Supplemental Fig. S9, C and D).

A short-term (24 h) labeling assay was performed with an isotope ^67^Zn to further investigate the role of OsbZIP48 on Zn uptake and distribution after Zn deficiency for 7 days (Fig. 5). Knocking out of *OsbZIP48* resulted in a lower Δ^67^Zn accumulation (net increased ^67^Zn during 24 h) in the shoots, and a higher Δ^67^Zn accumulation in the roots (Fig. 5A). However, the total amount of Δ^67^Zn accumulated in plants was no significant different between the *osbzip48* mutants and WT plants (Fig. 5B). The root-to-shoot translocation of Zn was much lower in the *osbzip48* mutants than in WT plants (Fig. 5C). These results suggested that the root-to-shoot translocation of Zn is impaired in the *osbzip48* mutants under Zn deficiency.

**Figure 5.**
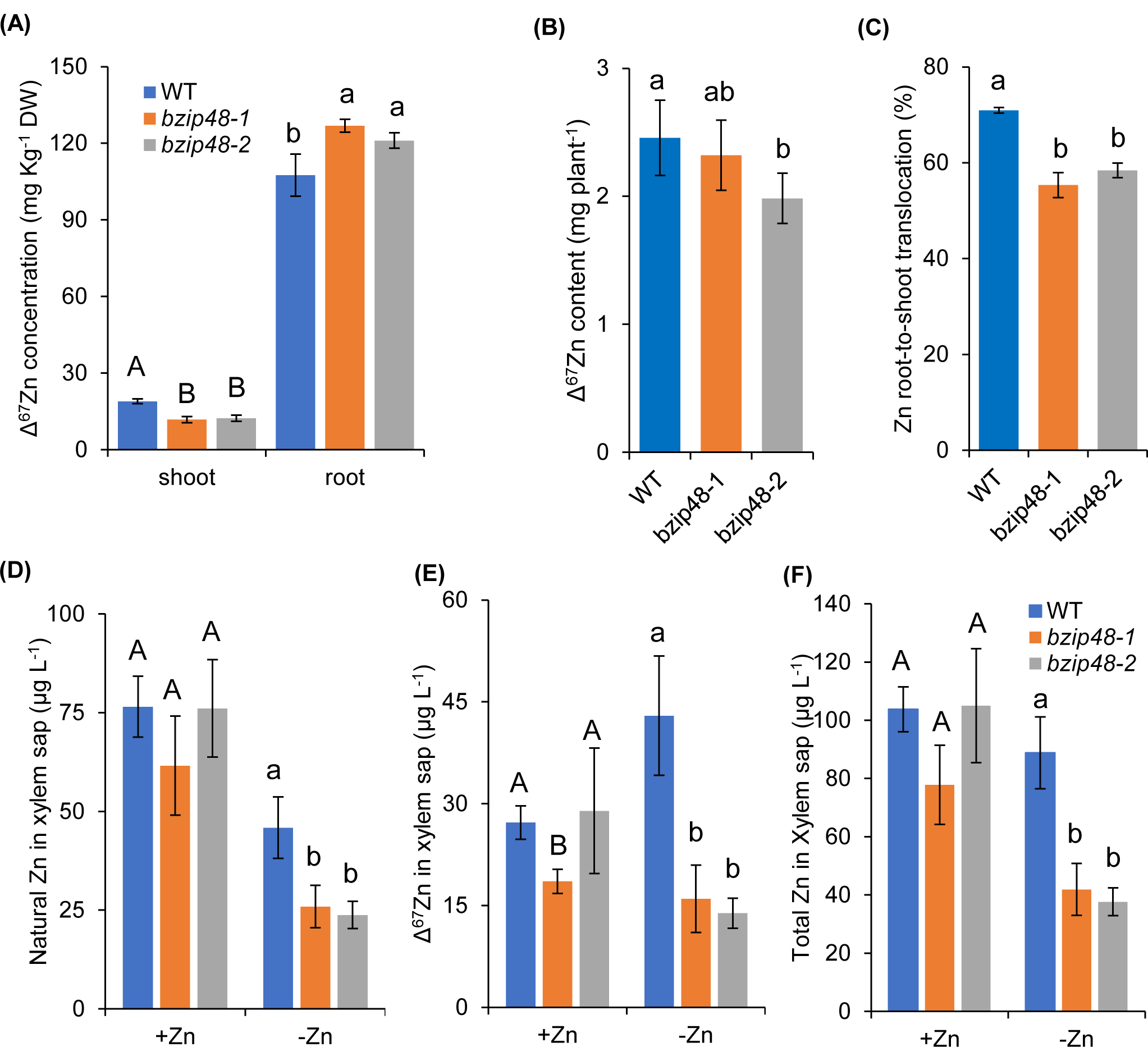
OsbZIP48 affects the root-to-shoot Zn translocation in rice under -Zn. A–C, short-term experiment with the stable isotope ^67^Zn. A, Δ^67^Zn concentration in shoots and roots. B, The total content of Δ^67^Zn per plant. C, ^67^Zn root-to-shoot translocation rate. Seedlings of WT and two *OsbZIP48* knockout lines were grown in a nutrient solution without Zn for 7 days and were then transferred to a nutrient solution containing 0.4 μM ^67^ZnCl_2_ for 24 hours. Shoots and roots were harvested separately and subjected to determination of ^67^Zn by ICP-MS. All data were based on comparisons with WT plants. DW: dry weight. Values represent means ± SD of biological replicates (n =4). (D–E) Natural Zn (D), net ^67^Zn (E), and total Zn (F) concentrations in xylem sap of WT plants and two *osbzip48* mutant lines. Four-week-old seedlings were transferred to nutrient solutions with or without Zn and, after 8 d of treatment, transferred to nutrient solutions with 0.4 μM ^67^ZnCl_2_ for 24 h. Xylem sap was collected at 10:00–11:00 am. Total Zn and ^67^Zn concentrations were measured by ICP-MS. Data are shown as mean ± SD values (n = 4). Significant differences between WT and *osbzip48* mutant lines were calculated by one-way ANOVA, followed by Tukey’s multiple comparison test. Different letters indicate significant differences (P<0.05).

Then, the Zn concentrations in xylem sap were analyzed. The concentrations of natural Zn, Δ^67^Zn, and total Zn (natural Zn + Δ^67^Zn) were determined. The natural Zn concentrations in xylem sap of rice were significantly decreased under Zn deficiency (Fig. 5D), to 60.0% (WT plants), 42.1% (*bzip48-*1), and 31.2% (*bzip48-*2) of their levels in plants under control conditions (Fig. 5D). The ΔZn concentration in xylem sap of WT plants increased significantly, and was about 0.58 times higher than that of the control (Fig. 5E). However, *bzip48-*1 and *bzip48-*2 mutants did not show similar changes, and only reached 86.2% and 47.9% of the control levels, respectively (Fig. 5E). Notably, the total Zn concentration in xylem sap of WT quickly recovered to that of the control after 24 h of ^67^ZnCl_2_ supplementation, while the total Zn concentration of *bzip48-*1 and *bzip48-*2 mutants reached only 53.8% and 35.8% of that of the control, respectively (Fig. 5F). These results further highlight the requirement of OsbZIP48 for upregulation of the root-to-shoot translocation of Zn in rice under Zn deficiency.

### OsbZIP48 plays an important role in global gene expression changes in -Zn conditions

Because of its role as a key transcriptional regulator involved in the response to Zn deficiency, the impact of OsbZIP48 on global gene expression was examined. Moreover, it was assessed whether the upregulated and downregulated genes in Zn deficiency treatment depended on functional OsbZIP48. These genes likely contain direct target genes of OsbZIP48. Therefore, transcriptome analysis was followed by functional validation of specific target genes.

RNA-seq obtained the transcriptome of WT and *osbzip48* mutant (*bzip48-*1) roots under control (+Zn) and Zn deficiency (-Zn) conditions for 7 d. A large number of differentially expressed genes (DEGs) were identified between WT grown under -Zn and +Zn conditions. *P* ≤ 0.01 and fold change (FC) ≥ 1.5 were used as cutoffs, and 979 genes (Cluster I) were identified to be induced and 1076 genes (Cluster II) were identified to be repressed by -Zn (Fig. 6A). The strong induction of *OsZIP4* in rice subjected to -Zn stress among this list confirmed the validity of the experimental design (Fig. 6B). Then, the functions of genes enriched within Cluster I genes and Cluster II genes were examined using Gene Ontology (GO) enrichment analysis. The most enriched GO enrichment biological process for Cluster I genes was “response to oxidative stress”, which was followed by “metal ion transport”, “transition metal ion transport”, “biological regulation”, “response to chemical stimulus”, and “regulation of primary metabolic process” (Supplemental Fig. S10A). However, “photosynthesis, light harvesting” was the most enriched GO enrichment biological process for Cluster II genes, which was followed by “photosynthesis”, “oxidation reduction”, “photosynthesis light reaction”, “metal ion transport”, and “peptide transport” (Supplemental Fig. S10B). These results indicated that genes involved in different biological functions were either induced or repressed in response to -Zn in rice roots.

**Figure 6.**
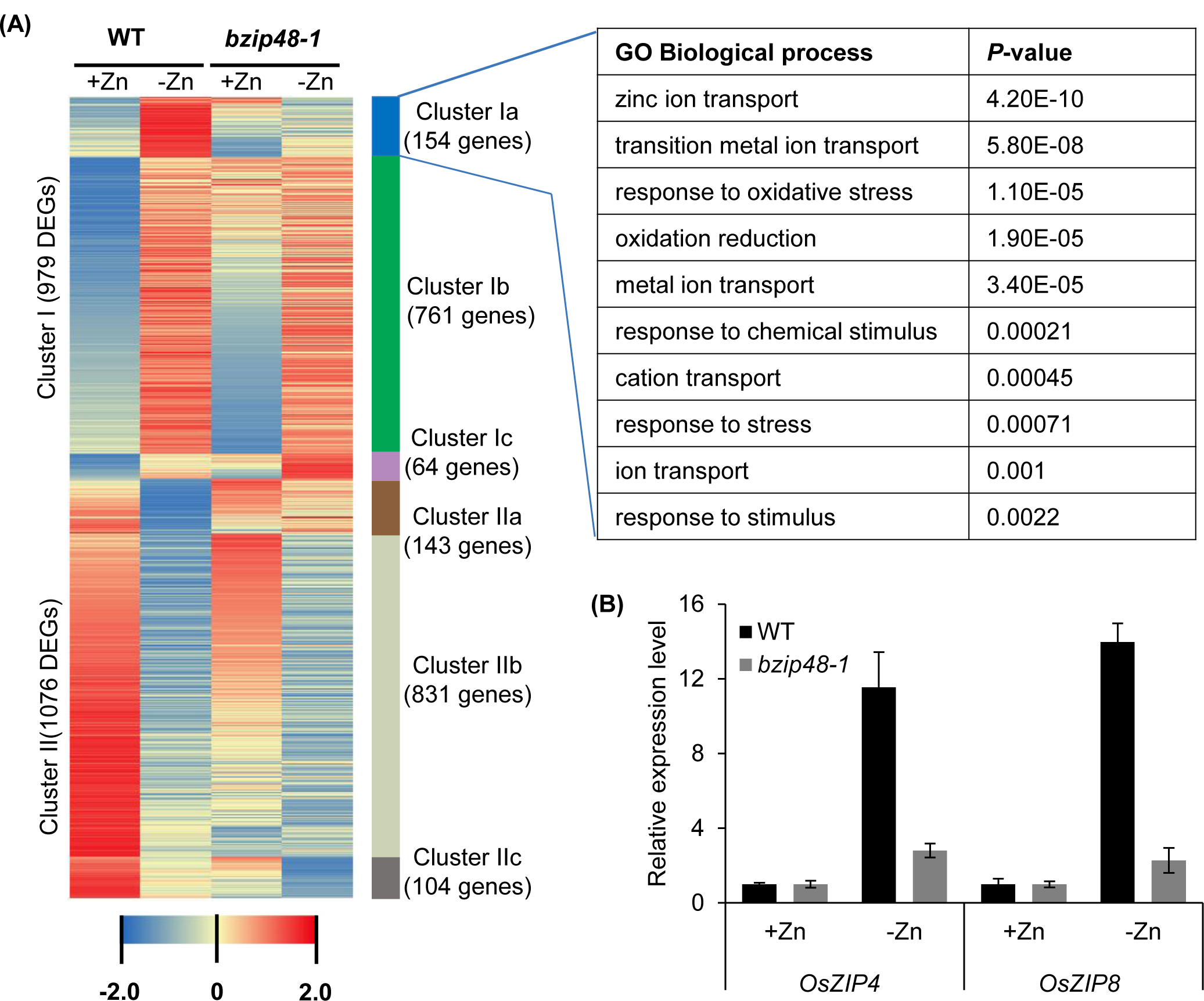
Zn deficiency-responsive genes are partially deregulated in the *osbzip48* mutant. (A) The 2055 Zn deficiency-responsive genes were hierarchically clustered and GO Enrichment (Biological process) was carried out for the genes in Cluster Ia. Genes with at least fold change (FC) ≥1.5 under -Zn conditions in WT roots are shown. (B) Relative expression levels of genes involved in Zn homeostasis in the *bzip48-1* mutant and WT plants, including *OsZIP4*, and *OsZIP8* were shown. Rice seedlings during the six-leaf stage were grown in solution with or without Zn for 7 d. Then, total RNA of root (B) were isolated for RT-qPCR analysis. Expression levels relative to WT plants for each gene under +Zn conditions were shown. Actin was used as the internal standard. Data are mean ± SD values (n = 3).

Next, it was explored how the loss of *OsbZIP48* affected the expression of Zn-regulated genes (Fig. 6A). Among the 979 Cluster I genes, 761 genes (Cluster Ib; e.g., *OsNAAT1* and *OsNAS3*) were normally regulated in *osbzip48* mutants (Fig. 6A and Supplemental Table S1). 64 genes (Cluster Ic; e.g., *OsHMA4* and *OsMT1G*) were hyperinduced in *osbzip48* mutants under -Zn conditions (Fig. 6A and Supplemental Table S2), suggesting that OsbZIP48 could either directly or indirectly function as a repressor of these genes. Moreover, 154 genes (Cluster Ia) were not induced (or induced to a lesser extent) in *osbzip48* mutants compared with WT under -Zn conditions (Fig. 6A and Supplemental Table S3). GO enrichment analysis showed that “Zn ion transport” was the most enriched GO enrichment biological process of Cluster Ia genes (Fig. 6A), including *OsZIP4* and *OsZIP8* (Supplemental Table S3). OsZIP4 and OsZIP8 have been implicated in Zn preferential distribution and/or translocation in rice (Ishimaru et al., 2005; Lee et al., 2010a; Lee et al., 2010b). These genes (involved in Zn translocation) were deregulated in *osbzip48* mutants, which is consistent with the Zn translocation analysis (Fig. 5, and Supplemental Fig. S9A, B). The GO categories “transition metal ion transport”, “response to oxidative stress”, “oxidation reduction”, “metal ion transport”, and “response to chemical stimulus” were also enriched (Fig. 6A). These results indicated that OsbZIP48 is either directly or indirectly involved in the activation of -Zn induced genes, including Zn-transporter genes and genes related to oxidative stress.

Among the 1076 genes that were repressed under -Zn conditions, 831 genes (Cluster IIb; e.g., *OsACS2*, *OsPSBR*, and *OsSWEET3A*) were normally regulated in *osbzip48* mutants (Fig. 6A and Supplemental Table S4). 104 genes (Cluster IIc; e.g., *OsABCG3* and *OsPR10A*) were hyperrepressed in *osbzip48* mutants under -Zn conditions (Fig. 6A and Supplemental Table S5), and 141 genes (e.g., *OsHAK13* and *OsbZIP27*) were not repressed (or repressed to a lesser extent) in *osbzip48* mutants compared with WT under -Zn conditions (Fig. 6A and Supplemental Table S6). This suggests that OsbZIP48 participates either directly or indirectly in repressing the expression of these genes involved in diverse biological processes.

Finally, to verify the potential direct regulation between OsbZIP48 and its potential target genes (mainly genes in Cluster Ia), 17 genes were selected that showed significantly lower expression in the *bzip48-1* line than in WT under -Zn conditions and were induced by -Zn (FC ≥ 4) in WT, for promoter analysis (Table 1). Most of these genes (14 out of 17) contained one or more ZDRE or ZDRE-like motif in their promoter sequences (Supplemental Table S7). *OsZIP4* and *OsZIP8* contained three ZDRE motifs and one ZDRE or ZDRE-like motif in their promoters, respectively (Supplemental Table S7). This suggests that OsbZIP48 also likely responds to -Zn in a ZDRE-dependent manner. To elucidate whether OsbZIP48 could directly activate the promoter of these candidate genes downstream, a luciferase (LUC) transient transcriptional activity assay was performed in tobacco leaves using the firefly LUC gene as a reporter, while OsbZIP48, driven by the CaMV 35S promoter, served as an effector (Supplemental Figure. S11A). Thus, 35S::OsbZIP48 co-expression with promoters of six genes (*pZIP4*::*Luc*, *pZIP8*::*Luc*, *pSFL1*::*Luc*, *pERF2*::*Luc*, *pPOX*::*Luc*, and *pDREB1B*::*Luc*) was observed to enhance fluorescent signal, and the expression level of LUC relative to control was obtained (Supplemental Figure. S11, B-E). This result suggests that OsbZIP48 could directly enhance the transcriptional activity of the promoters of these six genes. RT-qPCR analysis further confirmed that the upregulation of these genes under -Zn is OsbZIP48 dependent (Fig. 6B and Supplemental Fig. S12). These results suggest that OsbZIP48 regulates different pathways in response to -Zn, including the Zn transporter genes *OsZIP4*, and *OsZIP8*.

**Table 1.** Potential downstream genes of *OsbZIP48* in RNA-Seq analysis.

| TIGR locus ID | -Zn vs +Zn<br>(WT) |  | *bzip48-1* vs WT<br>(-Zn) |  | Description |
| --- | --- | --- | --- | --- | --- |
|  | Fold Change | FDR | Fold Change | FDR |  |
| LOC_Os04g21130 | 206.46 | 0 | 0.04 | 1.51E-11 | F-box protein PP2-B1 (***OsPP2B1***) |
| LOC_Os11g25040 | 89.72 | 7.74E-39 | 0.03 | 1.78E-08 | plant-specific domain family protein |
| LOC_Os06g47560 | 32.98 | 4.57E-02 | 0.10 | 2.50E-10 | expressed protein |
| LOC_Os10g18150 | 28.68 | 3.73E-39 | 0.05 | 1.18E-33 | splicing factor like 1 (***OsSFL1***) |
| LOC_Os03g09200 | 19.96 | 6.29E-11 | 0.11 | 1.70E-06 | domain of unknown function DUF966 |
| LOC_Os07g12890 | 13.85 | 1.98E-86 | 0.10 | 2.76E-120 | ***OsZIP8*** |
| LOC_Os04g47360 | 12.97 | 2.27E-231 | 0.14 | 1.76E-13 | OsPOP9 |
| LOC_Os08g10630 | 12.93 | 1.44E-97 | 0.16 | 3.82E-08 | ***OsZIP4*** |
| LOC_Os04g59210 | 11.00 | 2.02E-28 | 0.29 | 2.76E-14 | Peroxidase (***OsPOX***). |
| LOC_Os01g61840 | 9.25 | 3.35E-06 | 0.33 | 2.77E-03 | hypothetical protein. |
| LOC_Os06g46160 | 8.33 | 3.74E-71 | 0.30 | 1.25E-04 | expressed protein |
| LOC_Os07g11790 | 7.06 | 7.28E-03 | 0.07 | 1.05E-03 | Similar to terpene synthase 7. |
| LOC_Os03g64260 | 6.66 | 1.65E-13 | 0.31 | 4.07E-07 | ***OsERF2*** |
| LOC_Os09g01540 | 5.67 | 1.55E-02 | 0.23 | 3.08E-02 | Retrotransposon protein, putative |
| LOC_Os10g30150 | 5.49 | 1.42E-03 | 0.28 | 1.58E-03 | universal stress protein |
| LOC_Os06g01580 | 4.99 | 1.62E-07 | 0.48 | 1.36E-02 | Protease inhibitor |
| LOC_Os09g35010 | 4.83 | 1.47E-04 | 0.29 | 1.92E-03 | ***OsDREB1B*** |

Genes showing significantly lower expression in the *bzip48-1* line than WT under –Zn condition that were induced by –Zn (FC ≥ 4) in WT.

### OsbZIP48 positively regulates the expression of Zn transporter genes *OsZIP4* and *OsZIP8* by **directly binding to their promoters**

The binding of TFs to cis-regulatory elements in the promoter region of their target genes is a key step to control their expression, and they can act as activators or repressors. To determine whether OsbZIP48 binds to ZDRE motifs of the native promoter, one 43 bp region (*pZIP4*) surrounding the ZDRE motif in the *OsZIP4* promoter was used as bait for binding assays in the Y1H system (Fig. 7A). Interactions between OsbZIP48 and the promoter fragment were tested based on growth on media containing aureobasidin A. Yeast that was co-transformed with OsbZIP48 and the natural promoter regions (*pZIP4*) grew well on these selective media (Fig. 7A). Yeast co-transformed with OsbZIP48 and *pZIP4m* could not grow on these selective media, indicating that OsbZIP48 can bind to the ZDRE motif in the specific region of the *OsZIP4* promoter. Next, the above results were confirmed using EMSA, which identified the physical interaction of OsbZIP48 with the promoter sequence of *OsZIP4* and *OsZIP8* (Fig. 7B). Finally, ChIP-qPCR against *OsZIP4* and *OsZIP8* was used *in vivo* with the *35S::OsbZIP48-GFP* line (Supplemental Fig. S13). Three fragments, spanning different regions of the *OsZIP4* or *OsZIP8* promoters (*P1* to *P3*), were selected for RT-qPCR analysis (Fig. 7C). As shown in Fig. 7D, that fragments containing ZDREs (*P2* and *P3* for *OsZIP4*, and *P1* for *OsZIP8*) were considerably enriched compared with the negative control. These results demonstrated that OsbZIP48 binds to regions containing ZDRE motifs of *OsZIP4* and *OsZIP8* promoters, thus controlling their expressions in response to Zn deficiency.

**Figure 7.**
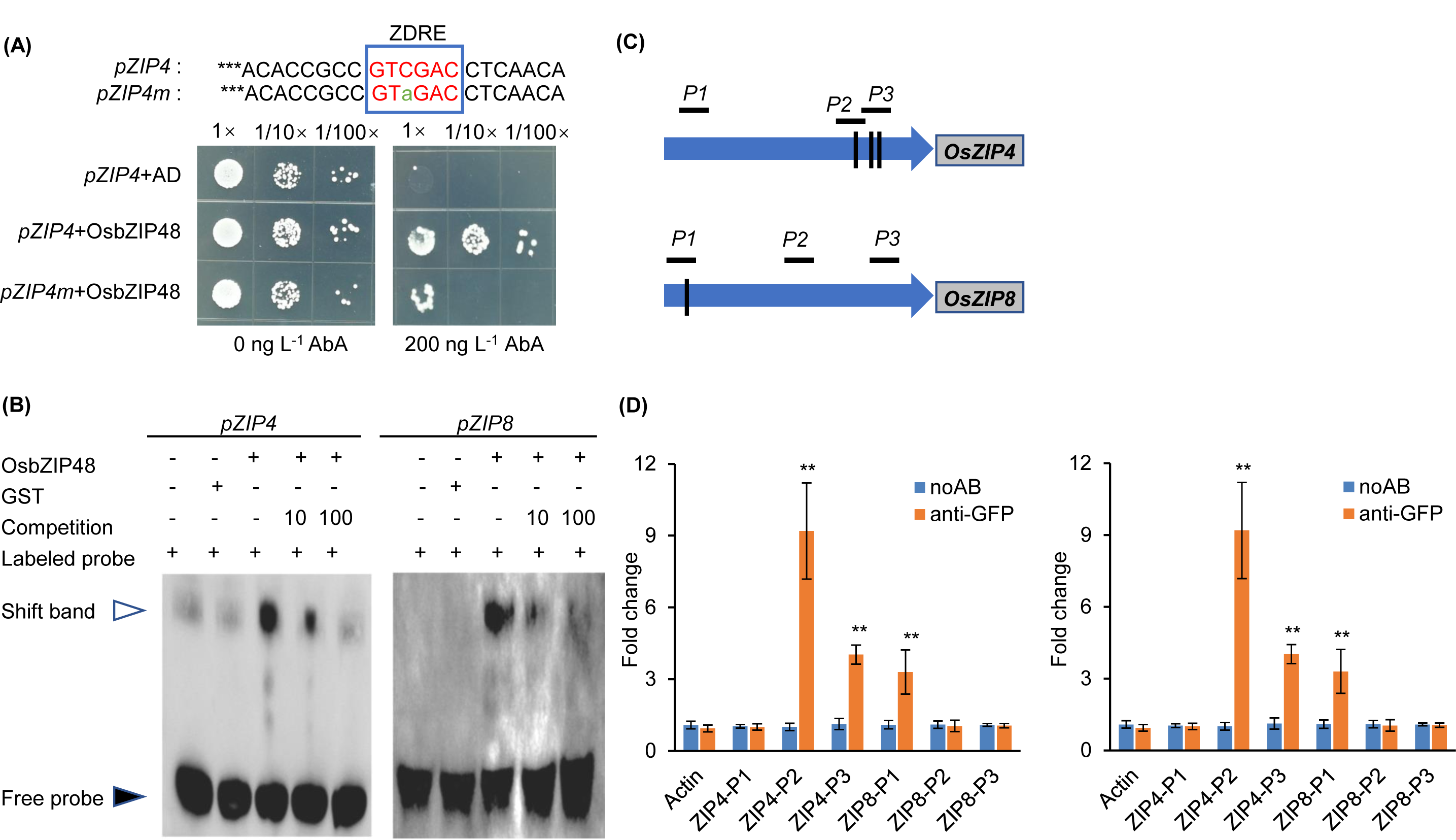
OsbZIP48 directly binds to the promoter of *OsZIP4* and *OsZIP8*. (A) OsbZIP48 binds to the *OsZIP4* promoter region containing the ZDRE motif in yeast. The small blue boxes within the promoter represent one ZDRE motif and *pZIP4m* indicates the mutated *pZIP4* (with a 1-bp mutation in the ZDRE motif). (B) Electrophoretic mobility shift assay (EMSA) results indicate that the interaction between recombinant OsbZIP48 and biotin-labeled probes containing the TGTCGAC motif in the *OsZIP4* and *OsZIP8* promoters. Arrows indicate the shifted bands and free probes. (C) Diagram of *OsZIP4* and *OsZIP8* promoters with three ZDRE motifs and one ZDRE motif, respectively. Black vertical bars represent ZDRE motifs. *P1*–*3* indicates genomic DNA fragments around *OsZIP4* and *OsZIP8* promoters for ChIP-qPCR. (D) ChIP-qPCR assay. Binding of OsbZIP48 to specific regions of the *OsZIP4* and *OsZIP8* promoters was examined with 4-week-old *35S*::*GFP-bZIP48* transgenic seedlings. The noAB (no antibody) and a fragment of *Actin* were used as negative controls. Data are means ± SD (n = 3). Statistical comparison between noAB and anti-GFP was performed by student *t*-test. Asterisks indicate significant differences from the control (\*\**P* < 0.01).

## Discussion

In this study, we demonstrated the essential role of the F-bZIP transcription factor, OsbZIP48 in Zn deficiency responses in rice. A recent study showed that OsbZIP48 could bind to the ZDRE motif and complete the zinc deficiency-hypersensitive Arabidopsis *bzip19bzip23* double mutant, suggesting it is a functional homolog of AtbZIP19 and AtbZIP23 in rice (Lilay et al., 2020). We showed that OsbZIP48 is localized in both nucleus and cytosol, and Zn deficiency significantly enhances its nuclear localization (Fig. 4, Supplemental Fig. S7), and it possesses transcription activator activity (Supplemental Fig. S2). We also showed that mutation of *OsbZIP48* significantly affects plant growth and development under Zn deficiency (Fig. 2 and 3, Supplemental Figure S4 and S5), these indicating that OsbZIP48 also functions as a transcription factor in rice essential for responses to zinc deficiency. However, different from *bZIP19* and *bZIP23*, the expression of *OsbZIP48* was not induced by Zn deficiency in roots but in shoots, and with a higher expression level in shoots (Assunção et al., 2010; Inaba et al., 2015; Fig.1). Furthermore, in *Arabidopsis*, mutation of one of the *bZIP* genes hardly affects plants, by contrast, the only *osbzip48* mutant is hypersensitive to Zn deficiency (Assunção et al., 2010; Fig.3, A-C). Double mutant of *bZIP19* and *bZIP23* mainly affected the uptake of Zn (Assunção et al., 2010) while knockout of *OsbZIP48* impaired the Zn translocation from root to shoot and Zn distribution to new leaves in rice under Zn-deficient conditions (Fig.5, Supplemental Figure S9). These findings indicate that OsbZIP48 is a major transcription factor of rice involved in Zn deficiency responses by regulating long-distance transport of Zn that effectively delivering Zn to new developing tissues required Zn utmost. Therefore, the growth impairment phenotype observed in the *osbzip48* mutant is not (or solely) due to low zinc uptake, but instead suggests a disturbance in zinc homeostasis as indicated by the RNA-seq analysis that will be discussed below. Whether the other two F-bZIP transcript factors, OsbZIP49 and OsbZIP50, are involved in Zn homeostasis regulation in rice requires further investigation (Lilay et al., 2020).

Early work in *Arabidopsis* identified the two essential TFs bZIP19 and bZIP23 for the physiological adaptation to Zn deficiency (Assunção et al., 2010; Inaba et al., 2015). Based on phylogenetic analysis and heterologous complementation assay on *Arabidopsis bzip19bzip23* double mutant, a number of their homologs (F-bZIP type TF) have been reported in different plant species, such as wheat (Evens et al., 2017), barley (Nazri et al., 2017) and rice (Lilay et al., 2020). Primarily, these TFs regulate target genes by recognizing the ZDRE or ZDRE-like motifs, and can partially rescue the growth phenotype of the *bzip19bzip23* double mutants under Zn deficiency, supporting a conserved mechanism of their roles in adapting to Zn limitation. On the other hand, although conserved in their Cys/His-rich motif and bZIP domain, which is characteristic of F-bZIP, the amino acids sequence of F-bZIPs from crops (barley, wheat, and rice) showed variations in their N-and C-terminal (Castro et al., 2017). Moreover, these F-bZIP TFs also showed different expression responses to Zn deficiency and complementation effect when expressed in the *bzip19bzip23* double mutant (Assunção et al., 2010; Inaba et al., 2015; Nazri et al., 2017; Lilay et al., 2020). These differences indicate that the transcriptional regulation mechanism of Zn-deficient responses is not completely the same between *Arabidopsis* and other plants. Clearly, the proper functional characterization of the proteins involved in the regulation of nutrient transport has to be ultimately validated in native species.

While the transcriptomic response to other microelements deficiencies (such as Fe deficiency) has been explored in detail, limited information on global gene expression in response to -Zn is available for common crops (Rodríguez-Celma et al., 2019). Such knowledge is of critical importance for the development of Zn-efficient or Zn-biofortified germplasms. The present study provides a genome-wide perspective of genes and biological pathways that respond to -Zn in rice at the early growth stage. Moreover, these results showed that among pathways that are regulated by the -Zn signal, the upregulation of GO categories such as “Zn ion transport”, “response to oxidative stress”, and “response to chemical stimulus” is OsbZIP48-dependent (Fig. 8). However, GO pathways such as “biological regulation” and “regulation of primary metabolic process” are not dependent on OsbZIP48 (Fig. 8). Therefore, this study proposes that the impairment in the regulation of Zn translocation, as well as the dysfunction of genes involved in the plant response to oxidative and chemical stimuli, is the underlying cause of the severe phenotype of *osbzip48* mutants under-Zn.

**Figure 8.**
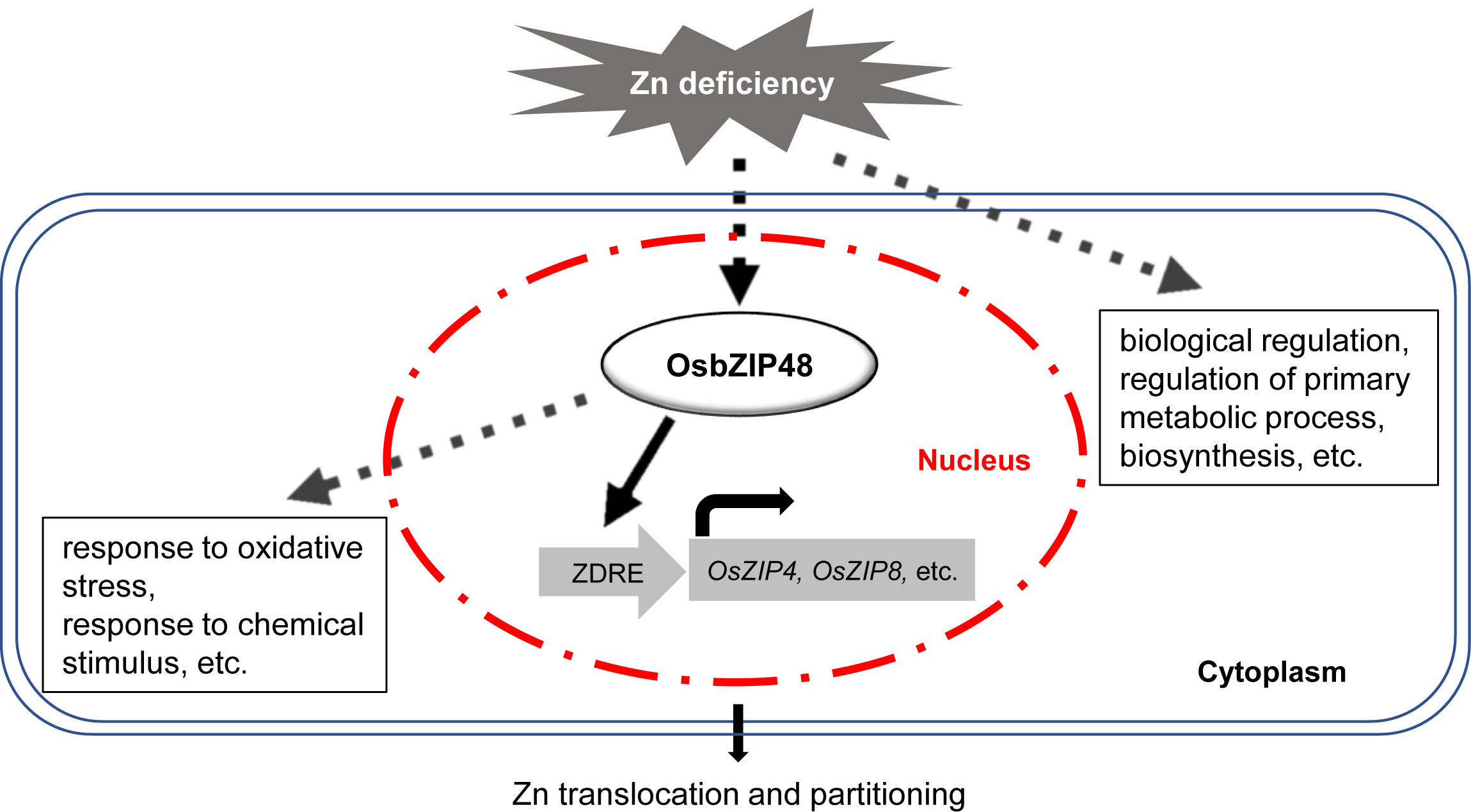
OsbZIP48-mediated regulation of Zn homeostasis in rice. In rice, Zn deficiency signal activates the downstream genes expression through an OsbZIP48-dependent and OsbZIP48-independent regulatory pathway. In the OsbZIP 48-dependent regulatory pathway, under Zn deficiency conditions, OsbZIP48 directly binds to the ZDRE motif in the promoter of -Zn responsive genes, including Zn transport related genes (*OsZIP4* and *OsZIP8*) that directly participate in Zn uptake, translocation and partitioning, and genes response to oxidative stress and chemical stimulus, etc. On the other hand, Zn deficiency signal also regulates other biological processes (eg. biological regulation, regulation of primary metabolic process, biosynthesis, etc.) that are not dependent on OsbZIP48. This Zn homeostasis regulation is essential for rice adapt to Zn fluctuations. In addition, other unknown factors are very likely working together with OsbZIP48 to perceive Zn deficiency signals and activates the downstream genes expression.

Furthermore, analysis of the obtained transcriptome led to the identification of potential targets for OsbZIP48. Among these genes, *OsZIP4* and *OsZIP8* were distinguished and were confirmed to show transcriptional and TF-DNA interactions (Fig. 6B, and Fig. 7; Supplemental Figure S11 B and D). Although *OsZIP10* was not identified in our RNA-seq analysis, it is also likely another candidate target of OsbZIP48 for containing three ZDRE motifs in its promoter (Lilay et al., 2020). Furthermore, the expression of *OsZIP5* and *OsZIP9*, two genes involved in Zn uptake, was upregulated under Zn deficiency conditions (Huang et al., 2020; Tan et al., 2020; Yang et al., 2020), however, whether OsbZIP48 plays a role in this process requires further investigation. Therefore, OsbZIP48 likely transmits -Zn signals in rice through Zn transporter genes such as *OsZIP4* and *OsZIP8* (Fig. 8). This is in line with previous work, which showed that overexpression of *OsZIP4* or *OsZIP8* led to excessive accumulation of Zn in roots and less Zn in shoots under Zn sufficient conditions in rice (Ishimaru et al., 2005; Lee et al., 2010b). Recently, our group demonstrated the critical role of OsZIP4 in delivering Zn to newly developing tissues through mode-based phloem transport (Mu et al., 2021). Consistently, the present study also showed that, in addition to a role in the root-to-shoot Zn translocation, OsbZIP48 also plays an important role in the transport of Zn to the developing tissues of rice under Zn deficiency (Fig. 5, and Supplemental Fig S9).

In addition to Zn ion transporters, there are several regulatory genes (such as *OsPP2B1, OsSFL1, OsERF2,* and *OsDREB1B*) downstream of OsbZIP48 (Table 1). *OsPP2B1* encoding F-box protein can selectively target proteins to ubiquitin-mediated protein degradation complex and plays an important regulatory role at the post-translational level (Maldonado-Calderón *et al*., 2012). Recently, many transcription factors involved in nutrient regulation were targeted by F-box proteins to gain a more complex regulatory network for nutrient homeostasis, such as the role of PRU1 in Pi homeostasis (Ye *et al*., 2018). *OsSFL1*, encoding a pre-mRNA splicing factor is also a potential target of OsbZIP48, splicing factor is involved in regulating mRNA splicing processes that is a post-transcriptional regulatory lay (Reddy et al., 2013). Interestingly one recent work showed that mutation in one splicing factor gene, *OsRS33*, significantly affects Zn translocation in rice, resulting in lower Zn in the shoot and higher Zn in the root (Dong *et al*., 2018). *OsERF2* and *OsDREB1B* encoded “Similar to Ethylene-responsive transcription factor 2” and “Dehydration-responsive element-binding (DREB) protein”, respectively, that both are phytohormone regulated transcription factor. However, their functions need to be identified on Zn homeostasis in further work.

In summary, this study provided evidence for the essential role of OsbZIP48 TF in maintaining Zn homeostasis as well as for the adaptation to fluctuations in the soil Zn status in rice. OsbZIP48 regulates key steps of Zn transport in rice by playing a critical role in controlling the root-to-shoot translocation of Zn under Zn deficiency (Fig. 8). This knowledge not only sheds light on the mechanism regulating Zn homeostasis in rice but also lays a solid foundation for biotechnological applications that enhance Zn uptake and accumulation in crops to contribute solving human nutrition problems.

## Materials and Methods

### Plant materials and growth conditions

Wild-type rice (*Oryza sativa* L. cv Nipponbare), two *osbzip48* mutants and GFP-tagged complementation lines with were used in this study. Rice seeds were soaked in water for 2 d before being transferred to a net floating on a solution containing 0.5 mM CaCl_2_. After 4 d, seedlings were transferred into half-strength Kimura B solution (pH 5.5) and cultivated in a greenhouse maintained at 25 to 30 °C. The composition of the nutrient solution was as follows: 0.18 mM (NH_4_)_2_SO_4_, 0.27 mM MgSO_4_·7H_2_O, 0.09 mM KNO_3_, 0.18 mM Ca(NO_3_)_2_·4H_2_O, 0.09 mM KH_2_PO_4_, 0.50 µM MnCl_2_·4H_2_O, 3.00 µM H_3_BO_3_, 1.00 µM (NH_4_)_6_Mo_7_O_24_·4H_2_O, 0.40 µM ZnSO_4_·7H_2_O, 0.20 µM CuSO_4_·5H_2_O, and 2.00 µM FeSO_4_. This solution was renewed every 2 d. Each physiological experiment was repeated at least three times with three replicates. The representative results are shown.

### Generation of *OsbZIP48* mutants and complementation lines

*osbzip48* mutants were generated by editing *OsbZIP48* in the ‘Nipponbare’ background using CRISPR/Cas9 system. Two OsbZIP48-specific target gRNA sequences (the primer sequences for gRNA1, 571–590 bp; gRNA2, 933–952 bp are shown in Supplemental Table S8) were inserted into the CRISPR/Cas9 vector pRGEB31, respectively (Xie & Yang, 2013). The resulting vectors were introduced into *Agrobacterium tumefaciens* strain EHA105 and *Agrobacterium*-mediated transformation was performed using calluses derived from mature rice embryos (Nishimura et al., 2006). The homozygous mutants were confirmed by sequencing the PCR products using primer pairs (primers are listed in Supplemental Table S8) flanking the two target sites, respectively. Two knockout lines of *OsbZIP48* (*bzip48-1* and *bzip48-2*) were selected for further phenotypic analysis as described below. The risk of off-target mutations was minimized by the selection of highly specific target sequences via a thorough genome search (www.genome.arizona.edu/crispr/).

To generate genetic complementary lines of *osbzip48* mutant, the genomic fragment of *OsbZIP48* containing the promoter (2137 bp region upstream of the start codon) and codon sequence (2956 bp genomic sequence from ‘ATG’) was amplified from rice leaf DNA, followed by subcloning fused with GFP into the binary vector pCAMBIA1300-sGFP. Primers used are found in Supplemental Table S8. The *ProbZIP48::bZIP48-GFP* vector was introduced into *osbzip48* mutant (*bzip48-1*) by *Agrobacterium*-mediated transformation rice transformation. Further GFP immunostaining and phenotype analysis on the complementation lines were described below.

### Gene expression analysis

To investigate the expression pattern of *OsbZIP48* at different growth stages, different tissues from plants (cv Nipponbare) grown in a paddy field from mid-June to the end of September were taken for RNA extraction. The soil Zn concentration were 134.12 mg kg^-1^, and soil pH was 5.6. The effects of deficiencies of essential minerals on *OsbZIP48* expression were investigated by exposing 2-week-old rice seedlings to nutrient solutions that either lack Zn, Fe, Cu, or Mn for 7 d; then, the roots and shoots were collected and separately subjected to RNA extraction. To examine the time-course of the expression pattern of *OsbZIP48* under -Zn, 2-week-old seedlings were exposed to +Zn or -Zn nutrient solution. Plant samples were harvested on days 1, 3, 5, 7, and 9 of the +Zn and -Zn treatment. The -Zn induced gene *OsZIP4* was used as a positive control.

Total RNA was extracted as described preciously (Hu et al., 2017) and used for first-strand cDNA synthesis with a HiScript II Q Select RT SuperMix (+gDNA wiper) kit (Vazyme, Nanjing, China) following the manufacturer’s instructions. Specific cDNA was amplified by Phanta^®^ Super-Fidelity DNA Polymerase (Vazyme), and RT-qPCR was performed using SYBR Premix Ex Taq™ (TaKaRa, Dalian, China) by a Mastercycler ep Realplex (Eppendorf). Three biological replicates were performed for each experiment. Normalized relative expression was calculated by the ΔΔC_t_ (cycle threshold) method, and *Actin* was used as the internal standard. The RT-qPCR primers used to amplify *OsbZIP48, OsZIP4* and *Actin* are found in Supplemental Table S8.

### Cellular and subcellular localization analysis

The *ProbZIP48::bZIP48-GFP* transgenic rice generated for complementation test of *osbzip48* mutant was used to examine the cellular localization of OsbZIP48. The immunostaining analysis was performed using an antibody against GFP following the procedure described previously (Yamaji and Ma, 2007). Longitudinal and cross sections (100 mm thickness) of root and basal stem were prepared using a microslicer (Linear Slicer PRO10; Dosaka EM). For cellular localization of OsbZIP48, GFP fluorescence from the sections of *ProbZIP48::bZIP48-GFP* transgenic seedlings were observed with a confocal laser scanning microscope (TCS SP8x; Leica Microsystems) according to Fu et al. 2019. The subcellular localization of OsbZIP48 was observed with a confocal laser scanning microscope (LSM710, Zeiss).

### Short-term experiment with the stable isotope ^67^Zn

The 7-day-old seedlings of WT and the knockout lines were grown in a nutrient solution without Zn for 7 days and were then transferred to a nutrient solution containing 0.4 μM ^67^ZnCl_2_ (97% enrichment; Taiyo Nippon Sanso) for 24 hours. Then shoots and roots were sampled separately for Zn determination. The roots were washed three times with 5 mM CaCl_2_ before harvest. Δ^67^Zn (net ^67^Zn increase) was calculated by Total ^67^Zn (Total Zn[^67^Zn]*4.1%) minus the natural abundance of ^67^Zn (Total Zn[^66^Zn]*4.1%). Concentration of ^66^Zn and ^67^Zn were measured by Inductively Coupled Plasma-Mass Spectrometry (ICP-MS, NexIon 300X; Perkin Elmer, Waltham, MA, USA) at isotope ^66^Zn (approximately 27.9% of total Zn in natural) and ^67^Zn (approximately 4.1% of total Zn in natural) mode. Three biological replicates were performed for each experiment. The translocation rate (%) of Δ^67^Zn was calculated by dividing its content in the shoot by its total content in the whole plant, which was the sum of Δ^67^Zn contents in all tissues.

### 67Zn content in xylem sap

The 4-week-old seedlings were transferred to nutrient solutions with or without Zn for 8 d, before transferring to a nutrient solution with 0.4 μM ^67^ZnCl_2_ for 24 h. For xylem sap collection, rice plants were cut to a height of 1 cm above their basal nodes. Xylem sap was collected with a pipette tip for 1 h. Four individual plants were pooled into for each sample. Xylem sap was collected at 10:00–11:00 am. All samples were centrifuged into new tubes, Δ^67^Zn and the natural abundance Zn were measured by ICP-MS. Total Zn is the sum of Δ^67^Zn and the natural abundance Zn. Three biological replicates were performed for each experiment.

### Metal concentration analysis

Rice tissues including shoot and roots, or leaves were harvested and washed three times with 5 mM CaCl_2_, twice in ultrapure water. Samples were dried at 85℃ for 3-5 days followed by digesting with 3 to 5 mL of HNO_3_/HClO_4_ (87:13, *v*/*v*) in a heating block at temperatures of up to 180℃ as described previously (Dong et al., 2018). Metal concentrations were determined by ICP-MS (PerkinElmer NexION 300X). For ^66^Zn and ^67^Zn determination, an isotope mode was applied.

### RNA-seq analysis

Two-week-old seedlings of *OsbZIP48* knockout line (*bzip48-1*) and wild-type rice were treated with (+Zn) or without (-Zn) 0.4 µM Zn for 7 d. Root samples (WT +Zn, WT -Zn, *osbzip48* +Zn, *osbzip48* -Zn) were harvested with three replications and rapidly frozen in liquid nitrogen. Totally 12 RNA samples were sent for RNA extraction as described above. The RNA quantity was determined with a ND-8000 spectrophotometer (Nanodrop Technologies, Inc., Wilmington, DE, USA), agarose gel electrophoresis, and a 2100-Bioanalyzer (Agilent Technologies, Santa Clara, CA, USA). RNA samples in the final analysis had been subjected to electrophoresis with no visible smears on agarose gels, with 260/280 ratios above 2.0 and RNA integrity numbers greater than 8.0. RNA samples were sent to Genewiz Biotechnology Corporation (http://www.genewiz.com.cn; Genewiz, Suzhou, China) for sequencing using the Illumina Hiseq2500 platform (Illumina, San Diego, CA, USA) according to the manufacturer’s instructions. A total of 48-95 M 101 bp paired-end reads were obtained for each sample. The reads were mapped to a merged reference gene GTF file of rice reference genome RAPDB (http://rapdb.dna.affrc.go.jp/download/archive/irgsp1/IRGSP-1.0_representative_2016-08-05.tar.gz, IRGSP-1.0_predicted_2016-08-05.tar.gz) and MSU 7.0 (ftp://ftp.plantbiology.msu.edu/pub/data/Eukaryotic_Projects/o_sativa/annotation_dbs/pseudomolecules/version_7.0/all.dir/all.gff3) databases (Dong et al., 2018) using TopHat (Trapnell et al., 2009). After normalization by estimating the dispersion factors in DESeq2 R package, a negative binomial generalized linear model fitting and a Wald significance test were then used to calculate the moderated estimation of fold change and *P*-values using DESeq2 (Love et al., 2014). Differentially expressed genes exhibiting 1.5-fold changes and Benjamini and Hochberg-adjusted *P*-values (FDR) ≤ 0.01 were selected. Significantly enriched GO terms and KEGG pathways compared with the genome-wide background were detected using the Hypergeometric test in our in-house pipeline with a threshold *P* value of ≤ 0.05 (Dong et al., 2018).

To verify the RNA-seq results, we performed RT-qPCR analysis of root RNA samples treated exactly the same as described above. We examined the expression of the following genes involved in Zn homeostasis: *OsZIP4* and *OsZIP8* (Ishimaru et al., 2005; Lee et al., 2010a; Lee et al., 2010b). Their specific primers are listed in Supplemental Table S8.

### ChIP-qPCR assay

A chromatin immuno-precipitation Kit (Merck Millipore, Burlington, MA, USA) was used to perform ChIP-qPCR assays. Briefly, 2g samples of 14-d-old 35S::bZIP48-GFP transgenic rice seedlings were fixed with 50 mL of 1.0% formaldehyde under vacuum for 10 min. Chromatin was extracted and sheared to 200 to 1000 bp fragments by ultrasonication according the instruction.. Then, 60 µL of sheared DNA was immune-precipitated with 4 mg of anti-GFP antibody (Santa Cruz Biotechnology, Dallas, TX, USA) overnight at 60 rpm and 4℃. DNA fragments that specifically associated with OsbZIP48 protein were released, purified, and used as templates for RT-qPCR. Genomic fragments from nonbinding sites were used as negative controls. Relative expression was calculated by the ΔΔC_t_ (cycle threshold) method, and *Actin* in “noAB” samples was used as the internal standard.

### Transactivation experiment in Yeast

To examine the transcriptional activation potential of OsbZIP48, the Open Reading Frame (ORF) of OsbZIP48 was amplified by PCR from rice root cDNA and subcloned in frame after the DNA binding domain of yeast GAL4 transcription factor (without activation domain) in pGBKT7 vector (pGBKT7-bZIP48). The pGBKT7-bZIP48 and control vector pGBKT7, were introduced into yeast strain AH109 that carried the GAL4-responsive HIS3 and LEU2 reporter gene together with pGADT7 vector and yeast cells were cultured on SD2 (-Trp-Leu) and SD4 (-Trp-Leu-His-Ade) media at 30℃ for 3–5 d according to the manufacturer’s manual (Clontech).

### Yeast one hybrid experiment

To investigate the interaction between OsbZIP48 protein and ZDRE motif of *OsZIP4* promoter, the Y1H assays were performed following the manufacturer’s instruction (Clontech Laboratories, Inc., Kusatsu, Japan). The promoter sequence with and without mutation in ZDRE motif were artificially synthesized and cloned into the *Kpn* I-linearized pAbAi vector (Clontech Laboratories, Inc., Kusatsu, Japan) to generate the bait vectors. The ORF of *OsbZIP48* from rice root cDNA, fused with GAL4 activation domain was amplified and subcloned into the prey vector pGADT7 (Clontech Laboratories, Inc., Kusatsu, Japan). Competent yeast cells were prepared according to the Clontech Yeast protocols handbook using the Y1H Gold yeast strain. The pAbAi vector harboring the ZDRE motif of *OsZIP4* promoter was integrated into the Y1H Gold genome. The level of auto-activation of the *pZIP4* fragment in yeast cells was examined on SD-Ura medium containing 200 µg L^-1^ aureobasidin A (AbA). Then, pGADT7-bZIP48 vector was transformed into yeast cells containing the *pZIP4* fragment and examined on SD-Leu medium containing 200 µg L^-1^ AbA. All the primers used for construction of Y1H vectors are listed in Supplemental Table S8.

### Electrophoretic mobility shift assay (EMSA)

For the expression analysis, the recombinant OsbZIP48 protein (GST-OsbZIP48), OsbZIP48 was amplified from ‘Nipponbare’ cDNA with GST-bZIP48-F and GST-bZIP48-R primers using KOD DNA polymerase (Toyobo, Osaka, Japan) and cloned into the pGEX-4T-1 vector using Exnase II (Vazyme Biotech Co., Ltd). The verified constructs were transformed into BL21 (DE3) competent cells, and transformants were cultured at 37℃ (until OD_600_ = 0.5) and then induced with 1 mM isopropylthio-β-galactoside (IPTG) for 16 hours at 23℃. The recombinant proteins were purified with MagneGST™ Protein Purification System (www.promega.com/protocols/; Promega, Madison, WI, USA).

DNA fragments of the *OsZIP4* and *OsZIP8* promoter containing the ZDRE motif were directly synthesized and 5’-end-labeled with biotin while unlabeled primers were used to produce competitors. Double-stranded oligo nucleotides were annealed by cooling from 95℃ to 37℃. EMSA assays were performed according to the manufacturer’s instructions for a LightShift Chemiluminescent EMSA Kit (Thermo Fisher Scientific, Waltham, MA, USA). The migration of the biotin-labeled probes was detected using ECL substrate (Thermo Fisher Scientific) and visualized using the ChemDoc XRS imaging system (Bio-Rad, Hercules, CA, USA). The probe sequences for EMSA are listed in Supplemental Table S8.

### Gene accession numbers

Gene accession numbers of all genes appearing in this text are listed in Supplemental Table S9. Illumina reads of all samples were deposited in the Sequence Read Archive at the National Center for Biotechnology Information (http://www.ncbi.nlm.nih.gov/sra) under accession number PRJNA643725.

## Funding information

This work was supported by the National Natural Science Foundation of China (32172668, 41907145 and 31770269); the National Key Research and Development Program of China (2021YFF1000400); the China Postdoctoral Science Foundation (2018M632317); funding from State Key Laboratory for Conservation and Utilization of Subtropical Agro-bioresources (SKLCUSA-b201709). The Plant, Soil, and Microbial Sciences Department at Michigan State University to H.R.

## Acknowledgments

The authors thank Prof. Jian Feng Ma (Okayama University, Japan) for critical reading and comments on the manuscript.

## Supplemental Data

**Supplemental Figure S1.** Expression pattern of Zn, Fe and Cu deficiency inducible genes in rice roots.

**Supplemental Figure S2.** Transcriptional activity of OsbZIP48.

**Supplemental Figure S3.** Schematic of CRISPR/Cas9 mutants of *osbzip48*.

**Supplemental Figure S4.** Rice seedling phenotypes at 15 d under -Zn.

**Supplemental Figure S5.** Leaf blade and sheath phenotypes of rice seedlings under -Zn.

**Supplemental Figure S6.** Rice seedling phenotypes under Fe, Mn, and Cu deficiency.

**Supplemental Figure S7.** Subcellular localization of OsbZIP48-GFP in *ProbZIP48*::*bZIP48-GFP* transgenic rice root cells.

**Supplemental Figure S8.** Zn content of WT and two *osbzip48* mutant lines under -Zn.

**Supplemental Figure S9.** OsbZIP48 affect the accumulation of Zn in young tissues under –Zn.

**Supplemental Figure S10.** Gene ontology (GO) abundance chart of the response genes to -Zn in WT roots.

**Supplemental Figure S11.** OsbZIP48 enhances the transcriptional of six downstream genes in tobacco leaves.

**Supplemental Figure S12.** Other genes could be regulated by OsbZIP48 in rice under -Zn. **Supplemental Figure S13.** Rice transgenic *35S*::*GFP-bZIP48* or *35S*:: *GFP* line for ChIP-qPCR assay were determined by immunoblotting analysis with anti-GFP antibody.

**Supplemental Table S1.** Genes normally expressed in *osbzip48* mutant under -Zn condition among -Zn-induced genes (761 genes).

**Supplemental Table S2.** Genes expressed higher in *osbzip48* mutant than WT under -Zn condition among -Zn-induced genes (64 genes).

**Supplemental Table S3.** Gene expressed lower in *osbzip48* mutant than WT under -Zn condition among -Zn-induced genes (154 genes).

**Supplemental Table S4.** Genes normally expressed in *osbzip48* mutant under -Zn condition among -Zn-repressed genes (831 genes).

**Supplemental Table S5.** Genes expressed lower in *osbzip48* mutant than WT under -Zn condition among -Zn-repressed genes (104 genes).

**Supplemental Table S6.** Genes expressed higher in *osbzip48* mutant than WT under -Zn condition among -Zn-repressed genes (141 genes).

**Supplemental Table S7.** ZDRE motifs in 17 possible downstream genes of OsbZIP48.

**Supplemental Table S8.** Primers used in this study.

**Supplemental Table S9.** Gene accession numbers are listed.

**Supplemental Table S10.** Genes induced by -Zn.

**Supplemental Table S11.** Genes repressed by -Zn.

## Notes

### Competing Interest Statement

The authors have declared no competing interest.

