## Supplementary material for "The transcription factor OsbZIP48 governs rice responses to zinc deficiency": Supplemental Figs.pdf

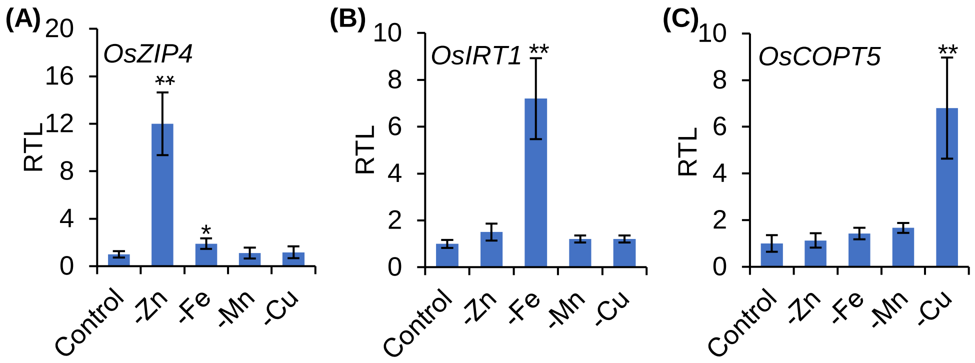


**Supplemental Figure S1.** Expression pattern of Zn, Fe and Cu deficiency inducible genes in rice roots. Response of *OsZIP4* (A), *OsIRT1* (B), and *OsCOPT5* (C) expression to metal deficiencies. Rice seedlings were grown in a nutrient solution without Zn, Fe, Mn, or Cu or with these trace elements (control) for 7 d. The expression levels of the three genes were determined by RT-qPCR. Expression levels relative to the control are shown. Actin was used as an internal control. Data are means ± SD of three biological replicates. Statistical comparison between control and treatment was performed by student *t*-test. Asterisks indicate significant differences from the control (*P < 0.01).

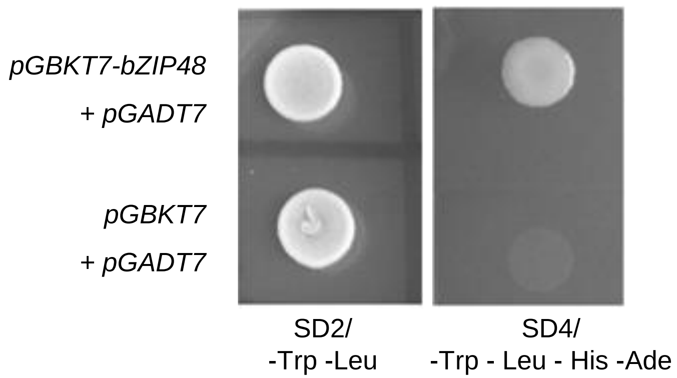


**Supplemental Figure S2.** Transcriptional activity of OsbZIP48.

The *OsbZIP48* open reading frame (ORF) was inserted between the *Eco*R I and *Bam*H I restriction sites of pGBKT7. Specific primers are shown in Table S1. The recombinant construct and the pGADT7 vector were co-transformed into yeast cells (strain, *AH109*). The transformed yeast cells were plated onto SD2 (-Trp-Leu) and SD4 (-Trp-Leu-His-Ade) media and incubated at 30 °C for 3–5 d to detect its growth.

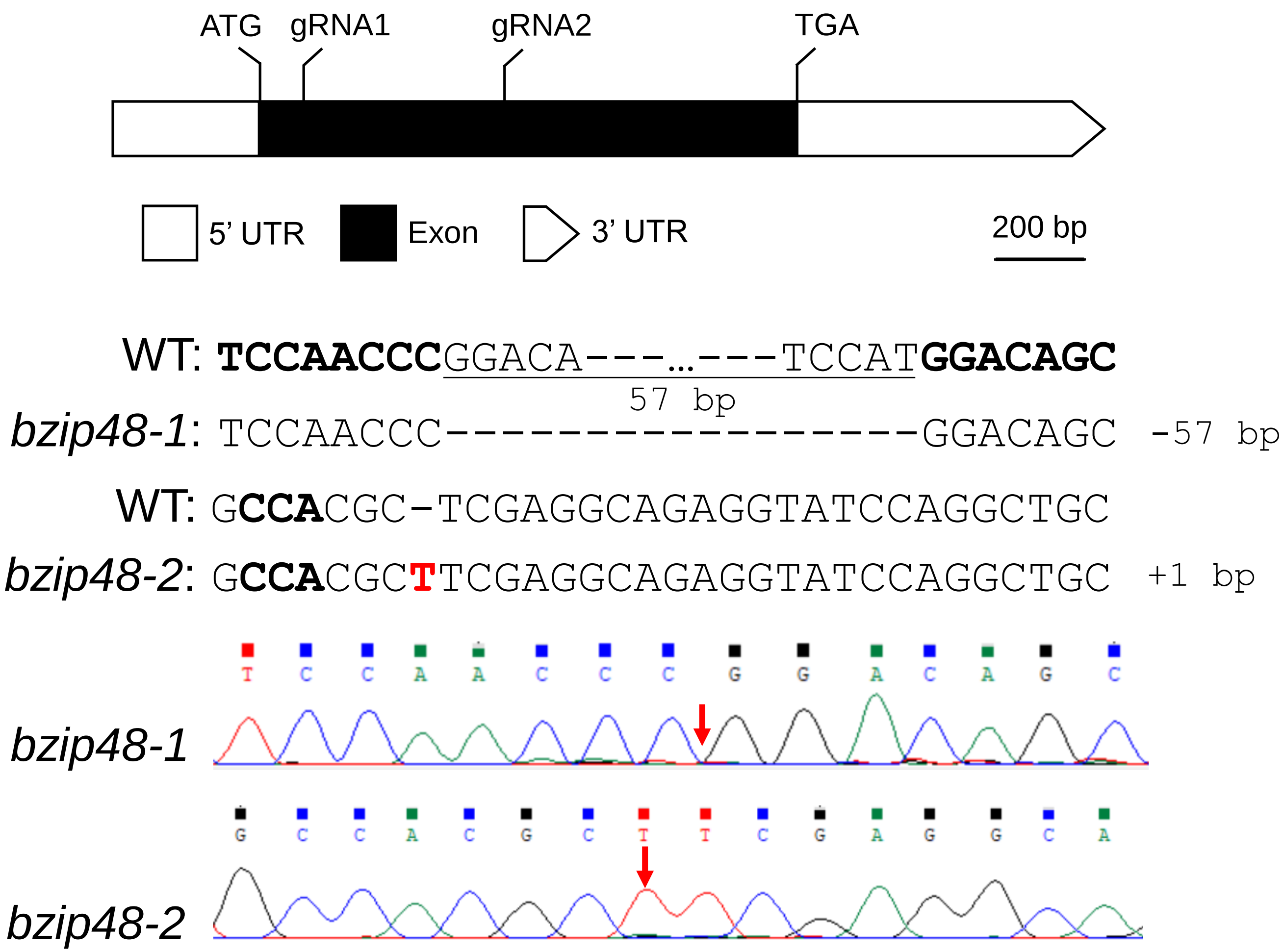


**Fig. S3** Schematic of CRISPR/Cas9 mutants of *OsbZIP48.*

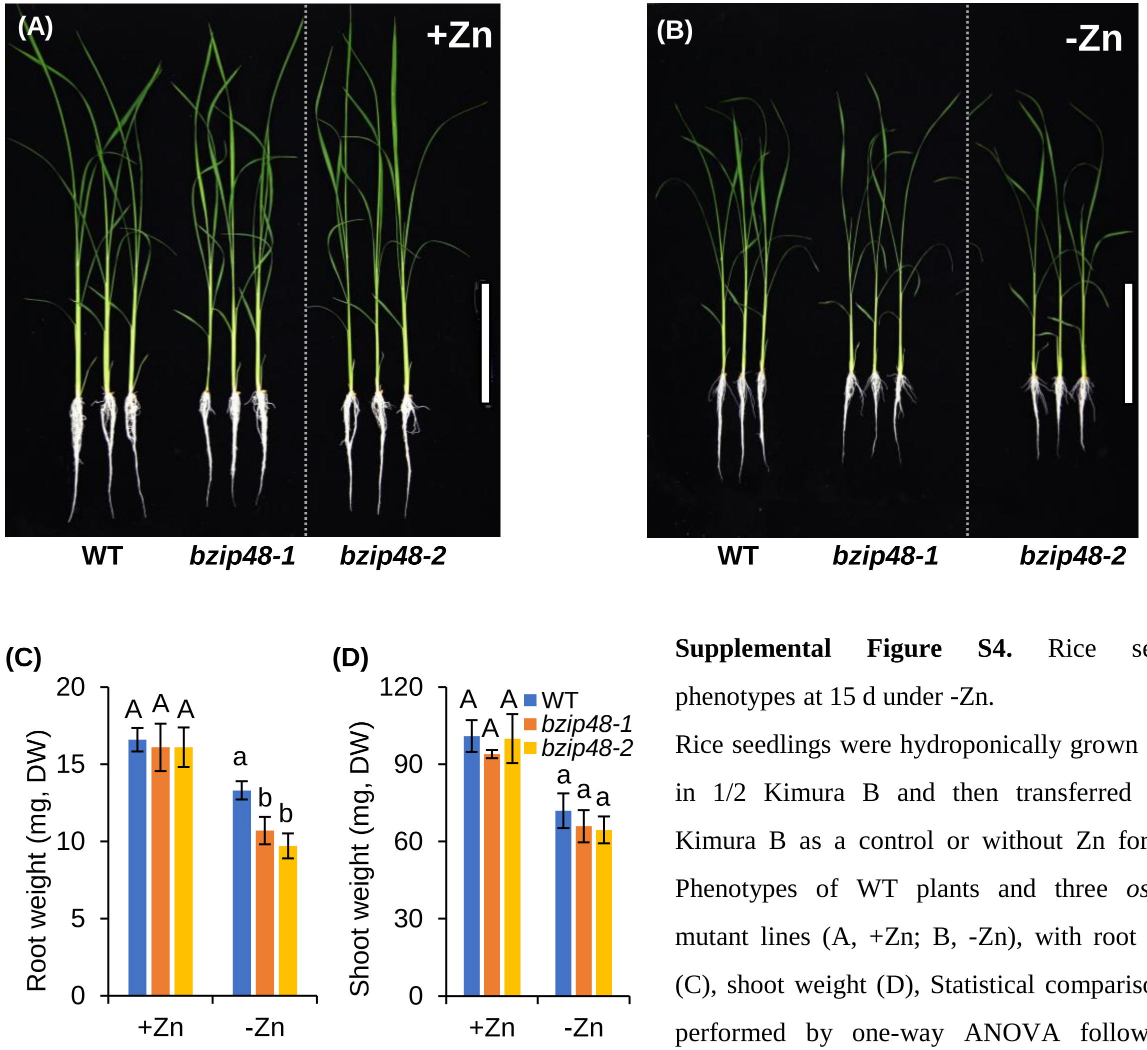


**Supplemental Figure S4.** Rice seedling phenotypes at 15 d under -Zn.

Rice seedlings were hydroponically grown for 6 d in 1/2 Kimura B and then transferred to 1/2 Kimura B as a control or without Zn for 15 d. Phenotypes of WT plants and three *osbzip48* mutant lines (A, +Zn; B, -Zn), with root weight (C), shoot weight (D), Statistical comparison was performed by one-way ANOVA followed by Tukey's multiple comparison test. Data are means ±SD (n = 5). Different letters indicate significant differences (P<0.05). Scale bars = 14 cm (A, B).

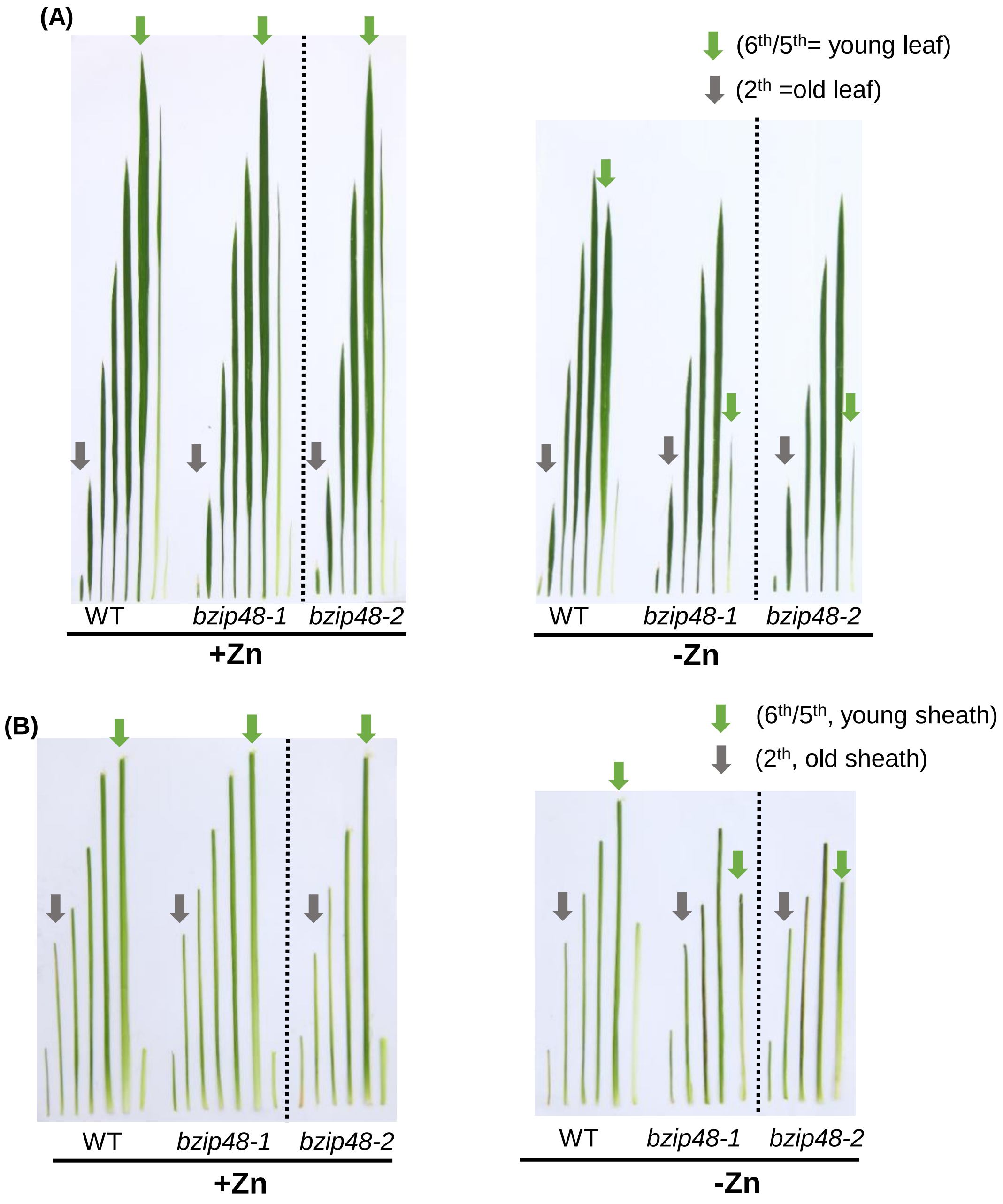


**Supplemental Figure S5.** Leaf blade and sheath phenotypes of rice seedlings at 15 d under -Zn.

The leaf blades (A) and sheaths (B) of each rice seedling grown in +Zn or -Zn solutions for 15 d are arranged chronologically, from oldest to youngest. The tissues indicated by these arrows were sampled and digested. Green arrows indicate a young leaf blade (A) and sheath (B), while gray arrows indicate an old leaf (A) and sheath (B).

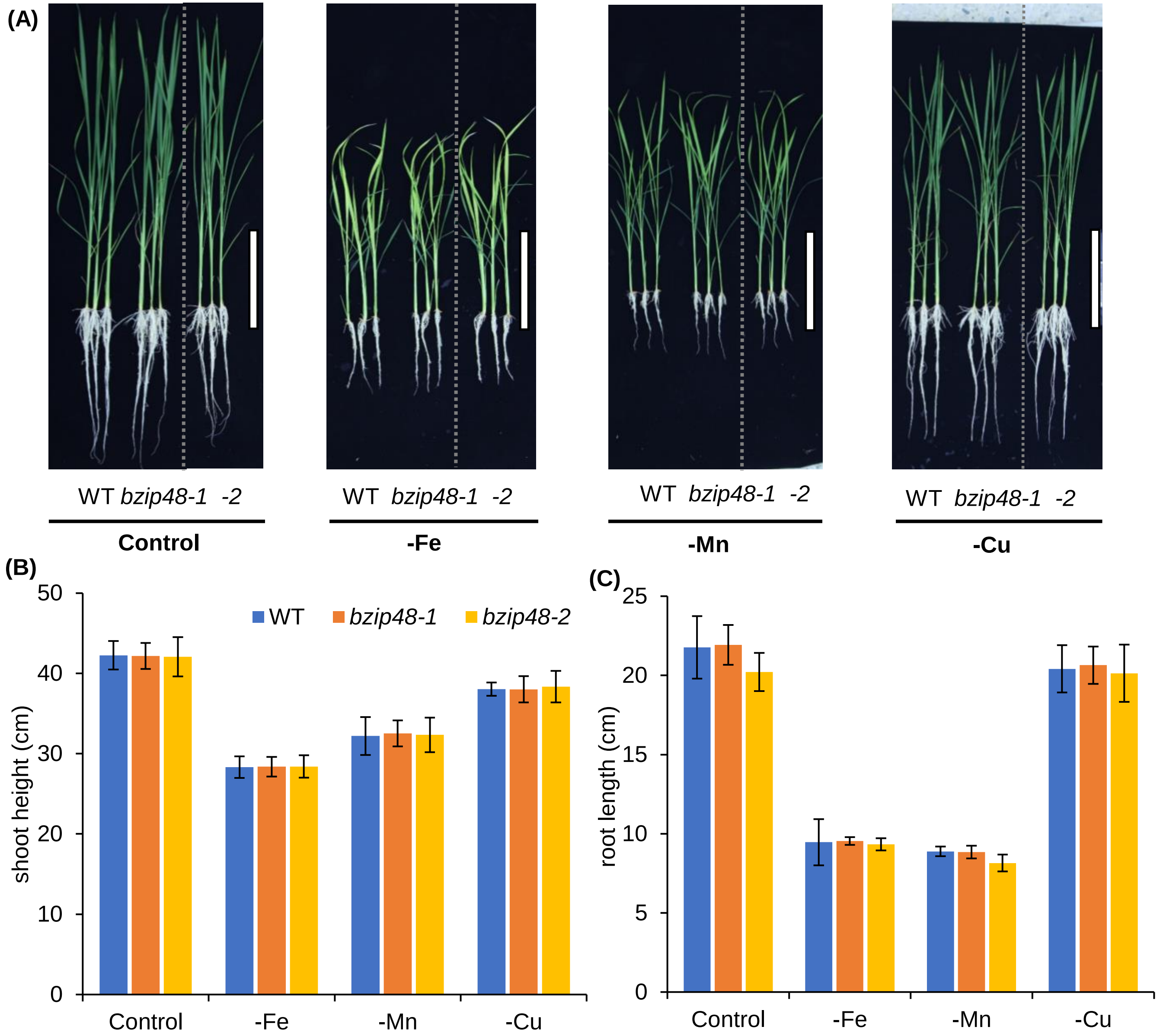


**Supplemental Figure S6.** Rice seedling phenotypes under Fe, Mn, and Cu deficiency for 25 d.

Rice seedlings were hydroponically grown for 6 d in 1/2 Kimura B and then in another solution (1/2 Kimura B or lacking Fe, Mn, or Cu) for 25 d. (A) Symptoms of element deficiencies appeared on the whole seedlings. Shoot height (B) and root length (C). Statistical comparison was performed by one-way ANOVA followed by Tukey's multiple comparison test. Data are means ±SD (n = 5). No significant differences were found between WT and mutant. Scale bars = 14 cm (A).

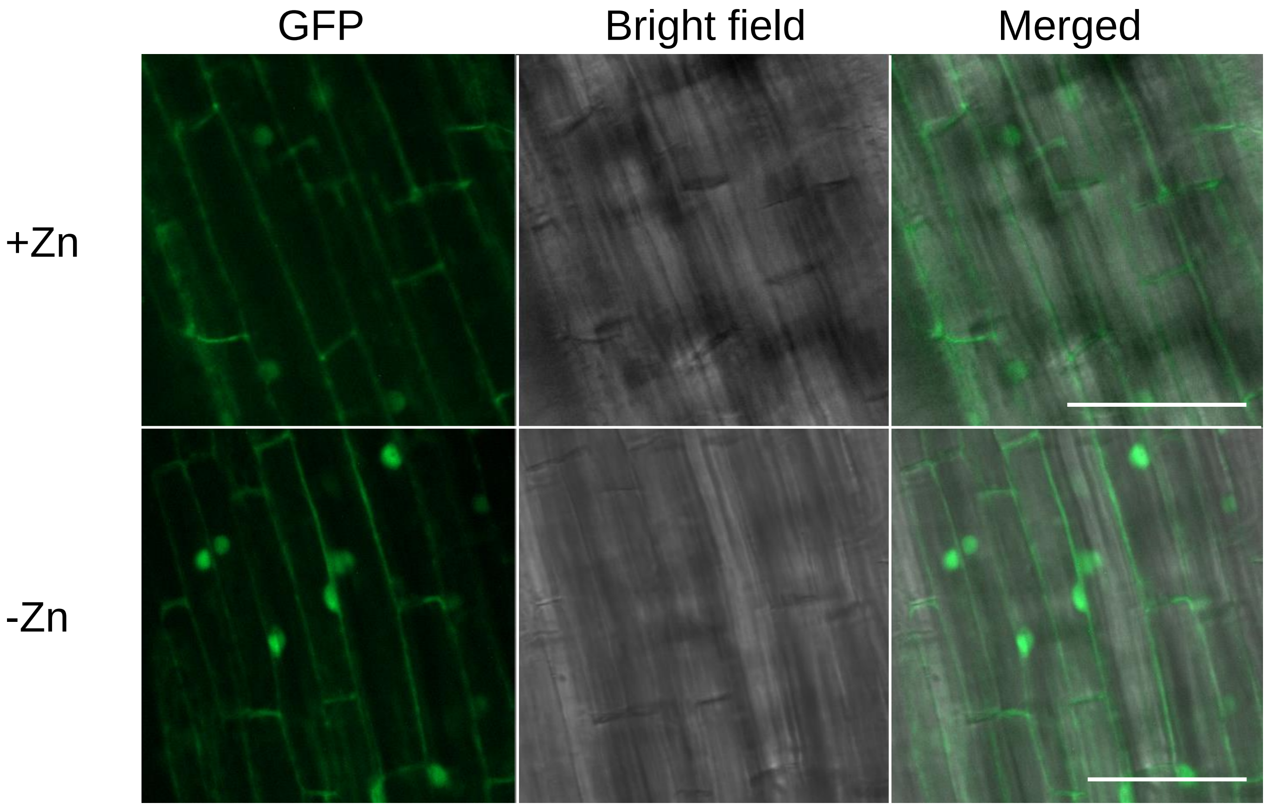


**Supplemental Figure S7.** Subcellular localization of OsbZIP48-GFP.

Subcellular localization of OsbZIP48-GFP in *ProbZIP48*::*bZIP48-GFP* transgenic rice root cells. Three-day-old rice seedlings were cultured in ½ Kimura B solution with (0.4 μM) or without (0 μM) Zn for 7 d, and cells of root elongation region were observed by laser confocal microscope. Scale bar = 50 μm.

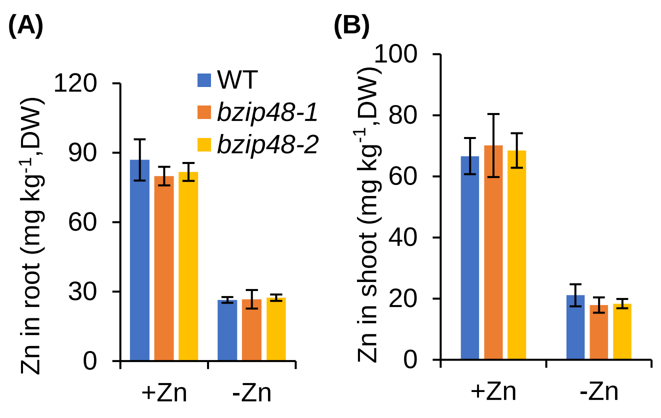


**Supplemental Figure S8.** Zn content of WT and three *osbzip48* mutant lines under -Zn for 15 d. Rice seedlings were hydroponically grown for 6 d in 1/2 Kimura B and then transferred to 1/2 Kimura B as a control or without Zn for 15 d. Zn concentration in root (A), and in shoot (B). Statistical comparison was performed by one-way ANOVA followed by Tukey's multiple comparison test. No significant differences were found between WT and mutant. Data are means ±SD (n = 5).

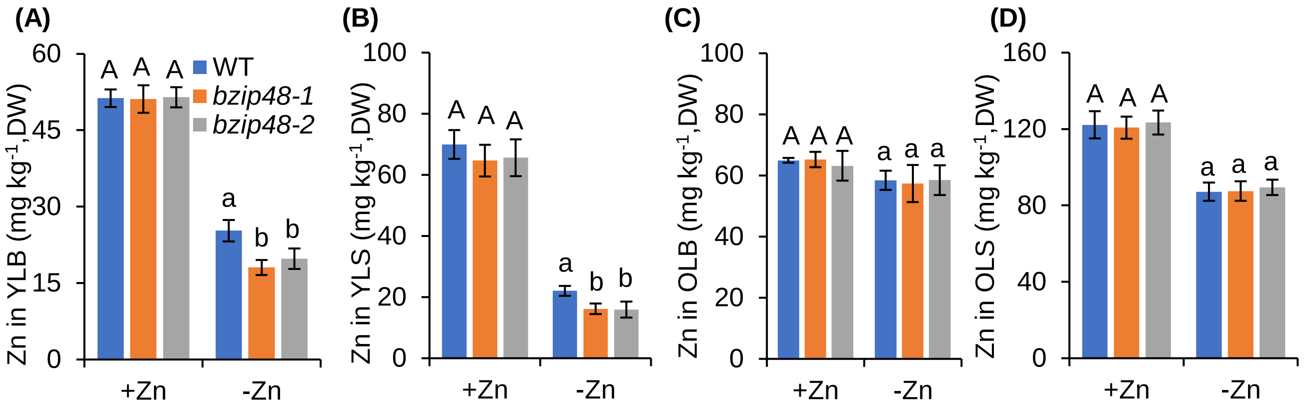


**Supplemental Figure S9.** OsbZIP48 affect the accumulation of Zn in young tissues under –Zn.

Zn concentration young leaf blades (YLB, A), young leaf sheaths (YLS, B) in old leaf blades (OLB, C), and old leaf sheaths (OLS, D), grown in +Zn or -Zn solutions were shown. Rice seedlings were hydroponically grown for 6 d in 1/2 Kimura B media and were then transferred to 1/2 Kimura B with 0.4 μM $ZnSO_4$ (+Zn) or without Zn (-Zn) for 15 d. Data are means ±SD (n = 3). Statistical comparison was performed by one-way ANOVA followed by Tukey's multiple comparison test. Different letters indicate significant differences (P<0.05). Data are means ±SD.

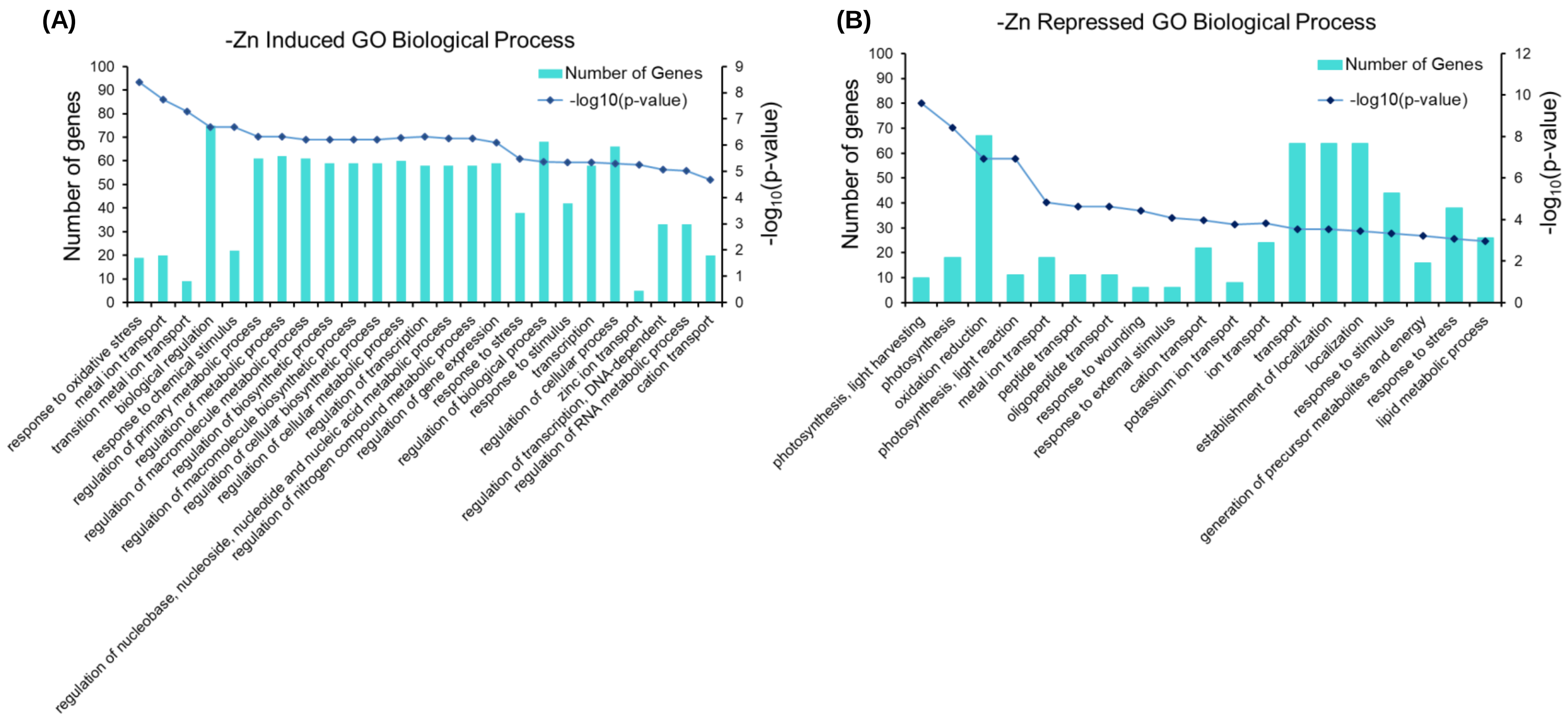


**Supplemental Figure S10.** Gene ontology (GO) abundance chart of the response genes to -Zn in WT roots.

(A) GO enrichment of -Zn induced genes (Cluster I; Supplemental Table S10); (B) GO enrichment of -Zn repressed genes (Cluster II; Supplemental Table S11).

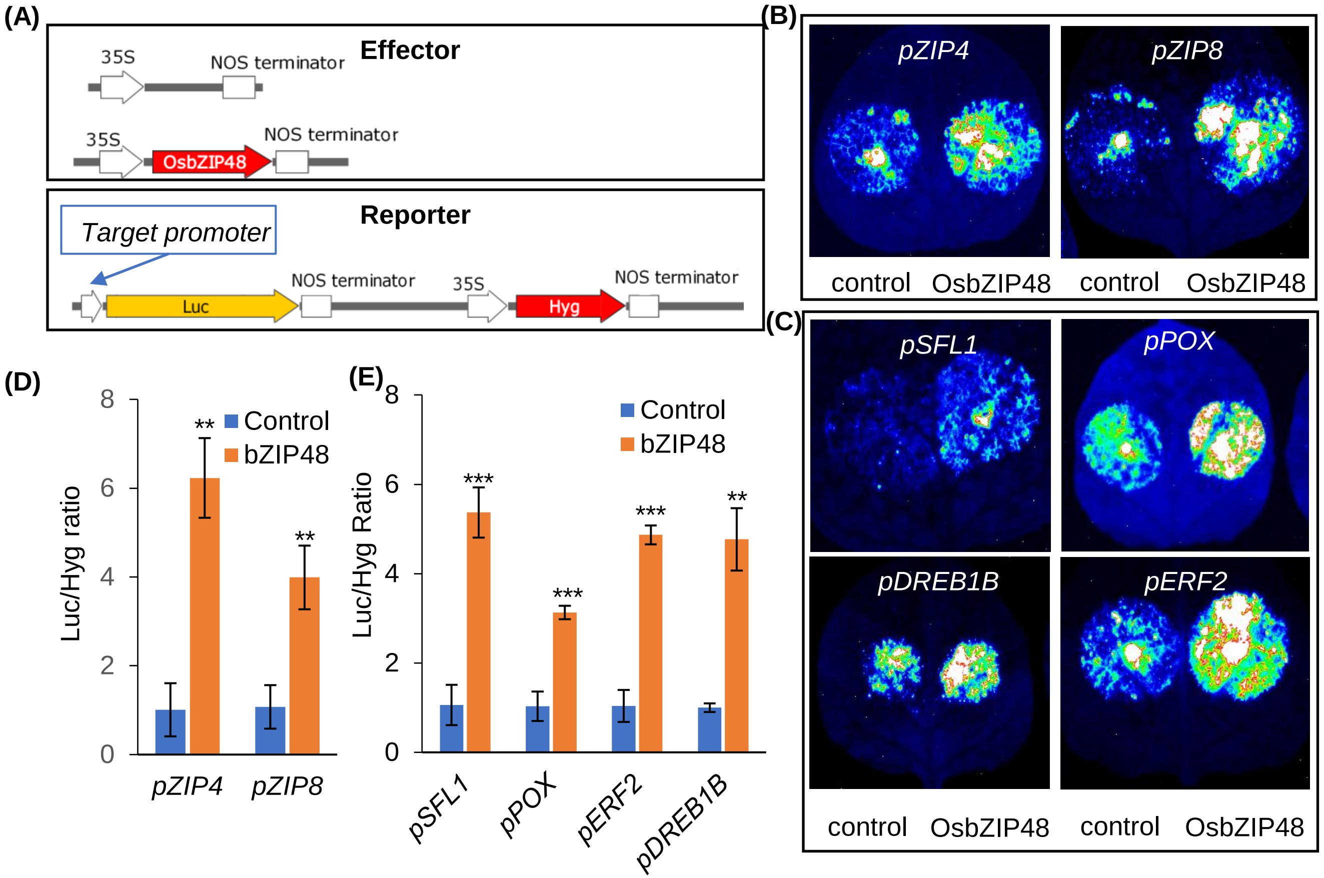


**Supplemental Figure S11.** OsbZIP48 enhances the transcriptional of six downstream genes in tobacco leaves.

(A) Schematic diagram depicting the constructs used in transient expression assays. (B) Luciferase signals were detected in leaf cells 48 h following co-infiltration with empty vector and pGWB502-bZIP48 and with p1381-*pZIP4::LUC* and p1381-*pZIP8::LUC*. (C) Relative LUC activity corresponds to the average of *pZIP4*::*LUC* and *pZIP8*::*LUC* observations across three independent replicates. Error bars represent the SD (n = 3). Asterisks indicate significant differences from the control (Student's *t*-test, $^{**}P < 0.01$, $^{**}P < 0.001$). (D) Transient luciferase reporter assays show the activation of *OsSFL1*, *OsPOX*, *OsERF2*, and *OsDREB1B* expression by OsbZIP48. Tobacco leaf transient expression assays using 1258-, 1396-,1397-, and 976-bp promoter fragments of *OsSFL1*, *OsPOX*, *OsERF2*, and *OsDREB1B*, respectively. (E) The relative expression levels of luciferase were quantitated following transfection with different vectors; *Hyg* was used for normalization (Student's *t*-test, $^{**}P < 0.01$ , $^{**}P < 0.001$).

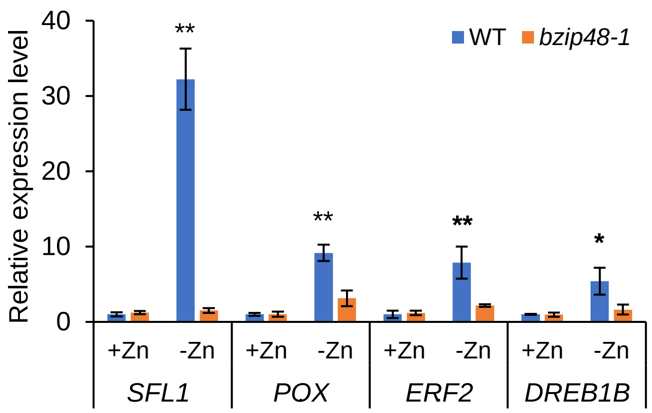


**Supplemental Figure S12.** Other genes could be regulated by OsbZIP48 in rice under -Zn. Quantitative analysis of relative expression level of other genes in the *bzip48-1* line and WT, including *OsSFL1*, *OsPOX*, *OsERF2* and *OsDREB1B*. Expression relative to each gene of WT under +Zn condition are shown. Rice seedlings during the six-leaf stage were grown in solution with or without Zn for 7 d, then total RNA of root was isolated for RT-qPCR analysis. *Actin* was used as the internal standard. Statistical comparison between WT and mutant was performed by student *t*-test. Asterisks indicate a significant difference from control ($^{*}P < 0.05$ and $^{**}P < 0.01$). Data are means ±SD (n=3).

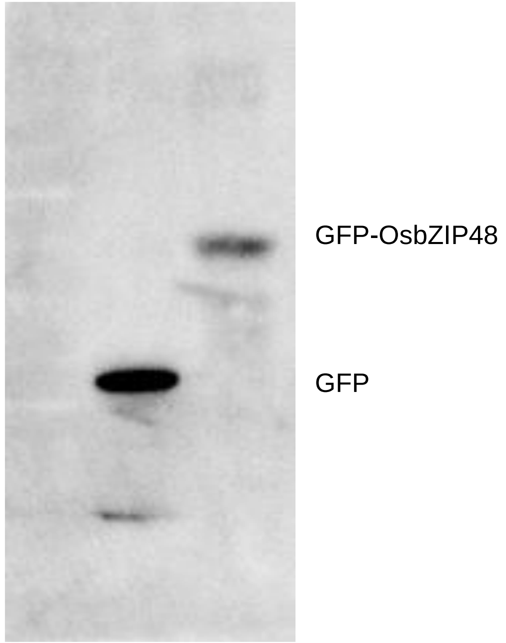


**Supplemental Figure S13.** Rice transgenic *35S*::*GFP-OsbZIP48* or *35S*:: *GFP* line for ChIP-qPCR assay were determined by immunoblotting analysis with anti-GFP antibody.
